# Behavioral state tunes mouse vision to ethological features through pupil dilation

**DOI:** 10.1101/2021.09.03.458870

**Authors:** Katrin Franke, Konstantin F. Willeke, Kayla Ponder, Mario Galdamez, Taliah Muhammad, Saumil Patel, Emmanouil Froudarakis, Jacob Reimer, Fabian Sinz, Andreas S. Tolias

## Abstract

Sensory processing changes with behavioral context to increase computational flexibility. In the visual system, active behavioral states enhance sensory responses but typically leave the preferred stimuli of neurons unchanged. Here we find that behavioral state does modulate stimulus selectivity in mouse visual cortex in the context of colored natural scenes. Using population imaging, behavior, pharmacology, and deep neural networks, we identified a shift of color selectivity towards ultraviolet stimuli exclusively caused by pupil dilation, resulting in a dynamic switch from rod to cone photoreceptors, extending their role beyond night and day vision. This facilitated the detection of ethological stimuli, such as aerial predators against the twilight sky. In contrast to previous studies that have used pupil dilation as an indirect measure of brain state, our results suggest that the brain uses pupil dilation itself to differentially recruit rods and cones on short timescales to tune visual representations to behavioral demands.

## Introduction

Neural responses are modulated by the animal’s behavioral and internal state, which allows to flexibly adjust information processing to guide behavior. This ubiquitous phenomenon is well-described across animal species, from invertebrates (Chiappe et al., 2010; Rowell, 1971) to primates (McAdams & Maunsell, 1999; Treue & Maunsell, 1996). In the mammalian visual cortex, neural activity is desynchronized during an active behavioral state (Reimer et al., 2014; Vinck et al., 2015; Zhuang et al., 2014), which can be reliably tracked by recording associated changes in pupil size (Reimer et al., 2014; Vinck et al., 2015) and locomotion activity (Niell & Stryker, 2010). The desynchronization is coupled with an enhancement of sensory responses and increases in signal-to-noise ratio (Bennett et al., 2013; Erisken et al., 2014; Niell & Stryker, 2010), resulting in better stimulus discriminability and detection (Bennett et al., 2013; Dadarlat & Stryker, 2017). Mechanistically, these effects have been linked to neuromodulators like actelycholine and norepinephrine (reviewed in Harris & Thiele, 2011; Lee & Dan, 2012) that act on different levels of the visual system. Other than changes in response gain, the tuning properties of visual neurons, like orientation selectivity or contrast preference, typically do not change across quiet and active states (Erisken et al., 2014; McAdams & Maunsell, 1999; Niell & Stryker, 2010). So far, however, this has largely been studied in non-ecological settings using synthetic stimuli like gratings.

In this work, we study how behavioral states modulate tuning properties of cortical neurons in mice in the context of naturalistic images. Critically, these scenes include the color domain of the visual input due to its ethological relevance: Across the animal kingdom, color vision is involved in a variety of distinct behaviors like mate selection, social communication, foraging, as well as predator and prey detection (reviewed in Gerl & Morris, 2008). Mice, like most mammals, are dichromatic and have two types of cone photoreceptors, expressing ultraviolet (UV)- and green-sensitive S- and M-opsin (Szél et al., 1992), respectively. Mouse rod photoreceptors homogeneously represent the visual scene. In contrast, UV- and green-sensitive cone photoreceptors predominantly sample upper and lower visual field, respectively, through an uneven distribution across the retina (Baden et al., 2013; Szél et al., 1992). Mouse UV-sensitive cones have been linked to predator detection (Baden et al., 2013; Qiu et al., 2021) – similar to other species (discussed in Cronin & Bok, 2016). In contrast, green-sensitive cones in mice are likely optimized for other tasks like prey hunting (Hoy et al., 2016; Johnson et al., 2021) and processing of coherent motion on the ground (Sit & Goard, 2020).

To systematically study the relationship between neural tuning and behavioral states, we combined *in-vivo* population calcium imaging of primary visual cortex (V1) in awake, head-fixed mice with deep neural network modeling. We extended a recently described neural network approach to predict population responses from naturalistic visual stimuli and the animal’s behavior (Sinz et al., 2018; Walker et al., 2019a). Compared to simpler models (Olshausen & Field, 2006), deep neural network models offer faithful predictions for cortical neurons in the visual system to arbitrary stimuli (Bashivan et al., 2019; Cadena et al., 2019; Walker et al., 2019a). Critically, this enabled us to characterize the relationship between neural tuning and behavioral states in extensive *in-silico* experiments without the need to control the behavior experimentally. From these *in-silico* experiments, we were able to derive specific predictions about neural tuning properties that we subsequently confirmed experimentally *in-vivo* (Bashivan et al., 2019; Walker et al., 2019a).

Using this approach, we found that color tuning of mouse V1 neurons systematically shifts towards higher UV-sensitivity during an active behavioral state. This shift was more pronounced for neurons sampling the upper visual field. By pharmacological manipulations of the pupil, we showed that this state-dependent change in color preference is exclusively caused by pupil dilation. This increases the amount of light entering the eye to an extent sufficient to result in a dynamic switch between rod- and cone-dominated vision, even for constant ambient light levels. We demonstrated that this change in spectral tuning selectively enhances discriminability of UV objects at the population level. Finally, we used synthetic stimuli inspired by natural scenes to show that the increased UV-sensitivity during active periods may tune the mouse visual system to enable better detection of overhead predators against the ultraviolet background of the sky.

Our results suggest that a dynamic shift from rod- to cone-dominated vision can be caused by rapid changes in pupil size across distinct behavioral states despite constant ambient light levels. As pupil dilation is also associated with brain states such as arousal and attention (Bradshaw, 1967; Hess & Polt, 1960), this bottom-up pupil mediated mechanism will interact with other top-down features of these states, such as neuromodulation, to rapidly tune the visual circuitry to changing behavioral requirements.

## Results

### Deep neural predictive models identify optimal colored stimuli of mouse V1 neurons

To study the relationship of neural tuning in mouse primary visual cortex (V1) and the animal’s behavior, we mainly focused on two well-described behavioral states in mice (Niell & Stryker, 2010; Reimer et al., 2014): A quiet state characterized by no locomotion and a small pupil and an active state indicated by locomotion and a larger pupil. We presented colored naturalistic images to awake, head-fixed mice positioned on a treadmill (Fig. 1a), while recording the population calcium activity of L2/3 neurons using two-photon imaging. To capture behavioral parameters of the animal, we simultaneously recorded locomotion activity, pupil size, as well as instantaneous change of pupil size (Fig. 1d, top), which all have been associated with distinct behavioral states (Niell & Stryker, 2010; Reimer et al., 2014). We back-projected visual stimuli on a custom light-transmitting Teflon screen using a DLP-based projector (Franke et al., 2019) with LEDs in the UV and green wavelength range (Fig. 1b). This allowed us to differentially activate UV- and green-sensitive mouse photoreceptors (see Methods). We presented 7,000 images from the ImageNet database (Deng et al., 2009), using the UV and green LED for the blue and green image channel, respectively. For the experiments, we selected images with high color contrast but similar mean intensity distributions across color channels (Suppl. Fig. 1). We recorded neural responses (from 500 to 1,000 neurons per recording) in V1 along its posterior-anterior axis (Fig. 1c; 650 x 650 μm at 15 Hz; Methods), sampling from various elevations across the visual field (Schuett et al., 2002). This choice was motivated by the gradient of spectral sensitivity of mouse cone photoreceptors across the retina, with higher UV-sensitivity in the upper visual field and higher green-sensitivity in the lower visual field (Baden et al., 2013; Szél et al., 1992). We presented each image for 500 ms, with interleaved gray periods lasting 300-500 ms (Fig. 1d, bottom).

**Fig. 1.**
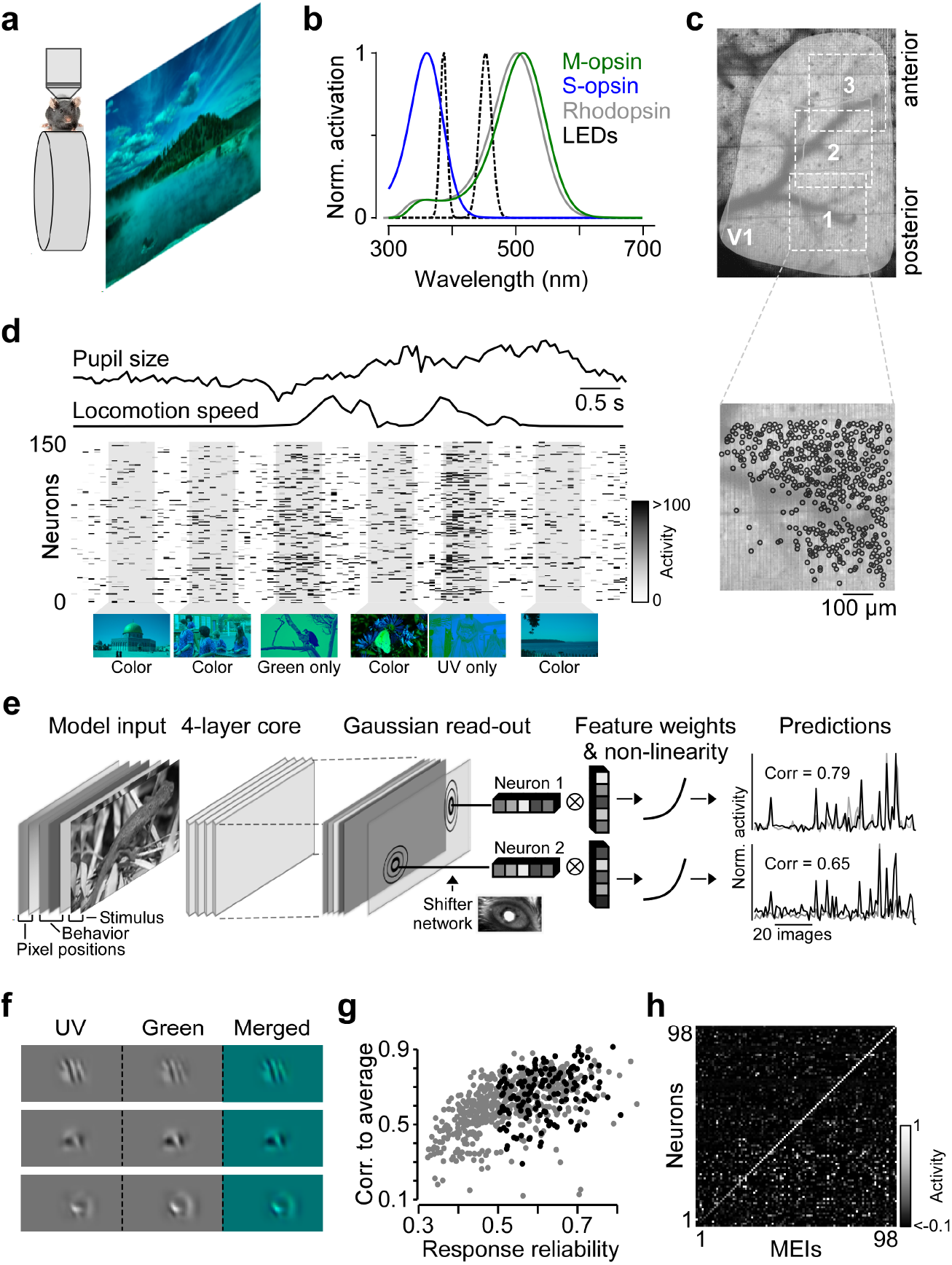
Experimental design, model and optimal colored images of mouse V1 neurons. **a**, Schematic illustrating experimental setup: Awake, head-fixed mice on a treadmill were presented with UV/green colored natural scenes (Methods and Suppl. Fig. 1). Stimuli were back-projected on a light-transmitting teflon screen using a DLP projector. **b**, Normalized sensitivity spectra of mouse short (S; blue) and medium (M; green) wavelength sensitive opsin expressed by mouse cone photoreceptors and rhodopsin (gray) expressed by rod photoreceptors, compared to LED spectra used for visual stimulation (dotted lines). **c**, Cortical surface through cranial window of a transgenic mouse expressing GCaMP6s in excitatory neurons, with positions of three scan fields (650 x 650 μm each) along the posterior-anterior axis of V1. V1 outline was identified based on reversals in retinotopic gradients (Garrett et al., 2014). Bottom image shows cells selected for further analysis (Methods). **d**, Activity of 150 neurons from (c, bottom) in response to colored natural scenes and simultaneously recorded behavioral data (pupil size and locomotion speed). Bottom images depict the presented images, with stimulus condition indicated below. **e**, Schematic illustrating model architecture: Model input consists of two image channels (UV and green), three behavior channels (pupil size, change in pupil size and locomotion speed) and two position channels encoding x and y pixel position of the input images ((Liu et al., 2018); see Methods and Suppl. Fig. 2). A 4-layer convolutional core with 64 feature channels per layer outputs a 3D tensor of feature activations for a given input. We then computed the neuronal responses by passing the core feature activations to a Gaussian readout and a subsequent non-linearity. The read-out position per neuron is learnt during training and adjusted according to recorded eye movement traces using a shifter network (Walker et al., 2019a). Traces on the right show average responses (gray) to 90 test images of two example neurons and corresponding model predictions (black). **f**, Maximally exciting images (MEIs) of three example neurons derived from the model. See Suppl. Fig. 3 for more examples. **g**, Response reliability to natural images (see Methods) plotted versus model prediction performance (correlation between prediction and average neural response to repeated presentations) of all cells of the exemplary scan. Neurons selected for experimental verification (inception loop) are indicated in black (see Methods). **(h**, Confusion matrix of inception loop experiment (Walker et al., 2019a), depicting each selected neuron’s activity to presented MEIs. Neurons are ordered based on response to their own MEI, from lowest (neuron 1) to highest (neuron 98). The majority of neurons (n=65) showed the strongest response to their own MEI.

We used a deep convolutional neural network (CNN) model to learn an *in-silico* representation of the recorded neuron population (Fig. 1e; Walker et al., 2019a), allowing us to systematically characterize the neurons’ tuning properties across behavioral states. We predicted neural responses as a function of both visual input and behavior by providing the model with the following input channels (Fig. 1e, left): (i) UV and green channel of the visual stimulus, (ii) three channels uniformly set to the recorded behavioral parameters (pupil size, instantaneous change of pupil size, and locomotion speed), and (iii) two channels encoding the x and y pixel positions of the stimulus image. We included the latter pixel-wise spatial location as input channels because it was previously shown to improve CNN model performance in situations when there are relations between high-level representations of the CNN and pixel positions in the image (Liu et al., 2018), such as a gradient in color across visual space. Our neural predictive models also included an additional network (shifter network; Walker et al., 2019a) that spatially shifts the receptive fields of all model neurons according to the recorded pupil position traces. For each dataset, we trained an ensemble of five 4-layer CNN models with a regression-based readout to predict the neuronal responses to individual images (Lurz et al., 2021). For all analyses, we used the average prediction across the ensemble (Hansen & Salamon, 1990). This yielded a correlation between response predictions and mean neural responses across repetitions of 0.56, and 0.546 ± 0.002 (mean ± s.e.m.) for the individual models (n=1,759 neurons from Fig. 2b; see also Suppl. Fig. 2). The fraction of variance explained (Pospisil & Bair, 2020) was 0.35 (0.314 ± 0.002 for individual models), which is comparable to the current best predictive models of mouse V1 (Lurz et al., 2021). We investigated the relative importance of the input and model components by performing a model comparison including simpler models. A model that utilized behavior, position encoding, and shifter network (default model) performed better at predicting neural responses than alternative models (Suppl. Fig. 2b).

**Fig. 2.**
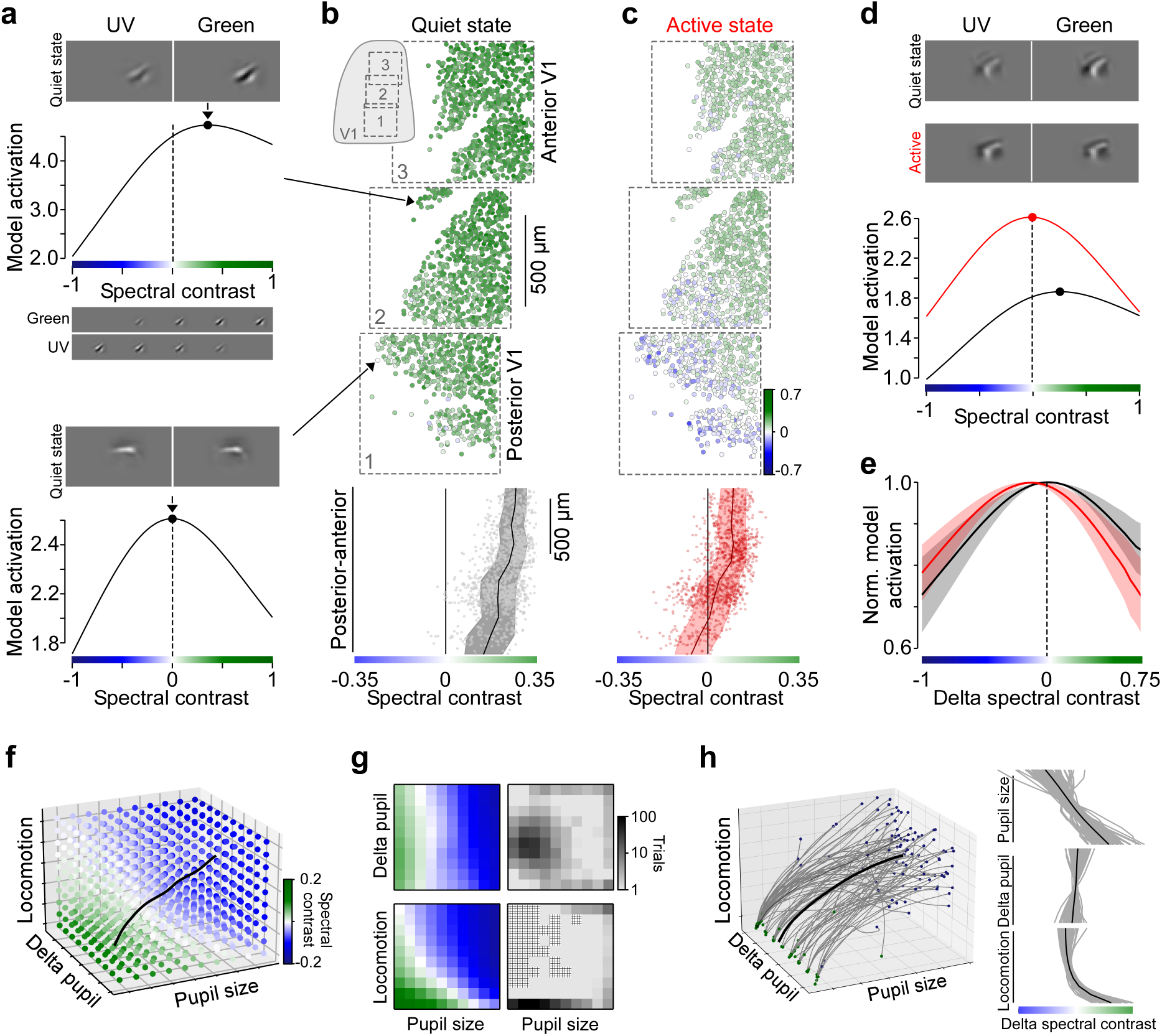
Color tuning of optimal colored images changes with behavioral state of the animal. **a**, MEI of an example model neuron optimized for a quiet state (3^rd^ percentile of pupil size and locomotion speed) and predicted model activation for varying spectral contrast levels (n=50). Example stimuli with distinct spectral contrasts plotted below - from all contrast in UV (left) to all contrast in green image channel (right). Bottom shows MEI and color tuning curve of another example model neuron. Dotted lines indicate spectral contrast of 0 and the MEI’s spectral contrast is marked on the peak of the curve. Arrows to the right indicate cortical position of selected neurons in (b). **b**, Neurons recorded in three consecutive experiments (n=1,759 neurons, n=3 scans, n=1 mouse) along the posterior-anterior axis of V1, color coded based on spectral contrast of their MEI optimized for a quiet state. Inset illustrates scan field positions within V1. Bottom shows MEI spectral contrast of all neurons from (b, top) along the posterior-anterior axis (dots) with binned average (black; n=10 bins) and s.d. shading (gray). Spectral contrast significantly varied across anterior-posterior axis of V1 (n=1,759, p<0.001 for smooth term on cortical position of Generalized Additive Model (GAM); see Methods and Suppl. Statistical Analysis). **c**, Like (b), but for MEIs optimized for an active state (97th percentile of both pupil size and locomotion speed). Spectral contrast significantly varied across anterior-posterior axis of V1 (n=1,759, p<0.001 for smooth term on cortical position of GAM; see Methods and Suppl. Statistical Analysis). **d**, MEIs of an example neuron optimized for quiet (black) and active (red) state, with corresponding color tuning curves below. **e**, Mean of peak-normalized color tuning curves of all neurons from (b, c), aligned with respect to peak position of quiet state tuning curves. Optimal spectral contrast was significantly shifted towards higher UV sensitivity during active (n=1,759, p<0.001 for behavioral state coefficient of GAM; see Methods and Suppl. Statistical Analysis), with only a small interaction between cortical position and behavioral state modulation (see Suppl. Statistical Analysis). **f**, Spectral contrast of MEIs optimized for n=1,000 different behavioral states, shown for an example neuron. Each dot represents a distinct state, color-coded based on spectral contrast of the corresponding MEI. Black curve indicates trajectory from behavioral state with highest to lowest spectral contrast. **g**, Spectral contrast of MEIs from (f), plotted separately for pupil size versus delta pupil (top) and pupil size versus locomotion speed (bottom). Right panels illustrate frequency of different behavioral states. Checkerboard pattern illustrates combinations of behavioral parameters not observed during the recording. **h**, Trajectories as shown in (g) for n=100 neurons from scan 1 in (b, c; gray) and mean trajectory (black). Start and end positions of trajectories (from highest to lowest spectral contrast) indicated by green and blue dots, respectively. Neurons were selected based on high prediction performance. Average difference in spectral contrast *±* s.d. of the trajectories for the three behaviors pupil size, delta pupil, and running speed were: 2.12 *±* 0.65 (Wilcoxon signed rank test: n = 100, p<0.001), −0.39 *±* 0.85 (Wilcoxon signed rank test: n=100, p<0.001), and 1.27 *±* 0.43 (Wilcoxon signed rank test: n=100, p<0.001) respectively.

Using our CNN ensemble model, we synthesized maximally exciting inputs (MEIs) for individual neurons *in-silico* (Fig. 1f; see Suppl. Fig. 3a for more examples). To this end, we optimized the UV and green color channels of a contrast constrained (Euclidean norm; see Methods) image to produce the highest activation in the model neuron using regularized gradient ascent (Methods; Bashivan et al., 2019; Walker et al., 2019a). We confirmed that computed MEIs indeed strongly drive the recorded neurons by performing inception loop experiments (Walker et al., 2019a). Specifically, after fitting the model with recorded responses and optimizing MEIs, we randomly selected MEIs of 150 neurons above a response reliability threshold for presentation on the next day (Fig. 1g). Visual responses to the same naturalistic images were highly consistent across days (Methods; Walker et al., 2019a). For most neurons (n=65 out of 98), the MEIs were indeed the most exciting stimuli: Neurons exhibited the strongest activation to their own MEI while showing little activity to the MEIs of other neurons (Fig. 1h). Together, this suggests that our modelling approach accurately captures tuning properties of mouse V1 neurons in the context of colored naturalistic scenes.

For the vast majority of neurons, MEI color channels were positively correlated (Suppl. Fig. 3a, b), suggesting that color-opponency, meaning that a neuron prefers an antagonistic stimulus in the UV and green channel, is rare for our stimulus paradigm. However, a fraction of neurons exhibited color-opponent temporal receptive fields in response to a full-field binary noise stimulus (Suppl. Fig. 3c-f) – in line with recent retinal work (Szatko et al., 2020). It is unlikely that the low number of color-opponent MEIs is due to an artifact of modelling, since our model faithfully recovered color preference and color-opponency of simulated neurons with Gabor receptive fields (Suppl. Fig. 4). This indicates that color-opponency of mouse V1 neurons depends on stimulus conditions, similar to neurons in mouse dLGN (Mouland et al., 2021).

### Color tuning of mouse V1 neurons changes with behavioral state

To study how color tuning of mouse V1 neurons changes with behavioral state, we performed a detailed *in-silico* characterization of the neurons’ color preference using the trained CNN model described above. We first focused on a quiet state with no locomotion and a small pupil (3^rd^ percentile of locomotion and pupil size across all trials) and an active state with locomotion and a larger pupil (97^th^ percentile of both locomotion speed and pupil size). For each neuron and distinct behavioral state, we generated a color tuning curve by predicting the neuron’s activity to varying color contrasts of its MEI (Fig. 2a).

Specifically, we systematically varied the ratio of contrast in UV and green MEI channel, while keeping the contrast across color channels constant. The spectral contrast of the modified MEIs ranged between −1 and 1 indicating extreme cases with all image contrast in the UV and green channel, respectively (see Methods). We then presented these modified MEIs to the model to obtain corresponding response predictions.

For both behavioral states, we found that the neurons’ optimal spectral contrast systematically varied along the anterior-posterior axis of V1 (Fig. 2a-c; for statistics, see figure legends and Suppl. Statistical Analysis): The UV-sensitivity significantly increased from anterior to posterior V1, in line with the distribution of cone photoreceptors across the retina (Baden et al., 2013; Szél et al., 1992) and previous work on cortex (Aihara et al., 2017; Rhim et al., 2017). For quiet behavioral periods, nearly all neurons (>99%) preferred a green-biased stimulus (Fig. 2b, bottom) – even the ones positioned in posterior V1 receiving input from the ventral retina, where cones are largely UV-sensitive (Baden et al., 2013). This observed distribution of color preferences across V1 indicates that visual responses in the quiet state are predominantly driven by rod photoreceptors which are green-sensitive, with only a small cone contribution (Rhim et al., 2021).

In contrast, during active periods we found that the color tuning of the neurons systematically shifted towards higher UV-sensitivity (Fig. 2c, d and Suppl. Fig. 6a-d). This was also accompanied by an overall increase in model activation (Fig. 2d), in line with previous results (Erisken et al., 2014; Niell & Stryker, 2010). The spatial structure of the MEIs was largely unchanged across behavioral states, with a pixel-wise correlation of *r* = 0.63 ± 0.24 across behavioral states (mean ± s.d., nN=1,759 neurons, Fig. 2d and Suppl. Fig. 5a-c). We additionally confirmed this by generating MEIs from two separate models, trained with trials from either the active or quiet state (Methods), resulting in similarly high correlations across behavioral states (*r* = 0.45 ± 0.33). The shift of color preference with an active state was depicted by a shift of the neurons’ color tuning curves towards more negative spectral contrast values indicating higher UV-sensitivity (Fig. 2e). It was slightly more pronounced for neurons in the posterior (−0.2 ± 0.05 change in spectral contrast) than the anterior V1 (−0.16 ± 0.05). For active behavioral periods, neurons in posterior V1 exhibited UV-biased MEIs, while neurons in anterior V1 largely maintained their preference for green-biased stimuli, consistent with a cortical distribution of color tuning expected from cone-dominated visual responses (Rhim et al., 2021).

When extending our *in-silico* analysis beyond these two extreme behavioral states to intermediate behavioral parameters, we found that color preference predominantly varied with locomotion speed and pupil size, while rapid changes in pupil size associated with internal states (Reimer et al., 2014) played a minor role in explaining the neurons’ color preference (Delta pupil, Fig. 2f-h). To this end, we systematically investigated the relationship between V1 color tuning and the animal’s behavior by sampling 1,000 combinations along the recorded behavioral parameters and optimized MEIs for each state. Note that some combinations of behavioral parameters do not exist in the data (Fig. 2g) because pupil size and locomotion speed are highly correlated. For a single neuron, color preference smoothly varied in this three-dimensional behavioral space, illustrated by the trajectory from highest to lowest spectral contrast value (Fig. 2f). The trajectories of the 100 best predicted neurons from scan 1 in Fig. 2b showed that color preference consistently changes with both increases in pupil size and locomotion speed (Fig. 2h). Interestingly, spectral contrast linearly varied with pupil size, but was non-linearly modulated by locomotion speed. The latter might be due to the fact that there are two rather discrete states of locomotion speed (Fig. 2g, bottom right).

We confirmed the above prediction from our *in-silico* analysis that mouse V1 color tuning shifts towards higher UV-sensitivity during active periods using a well established sparse noise paradigm for receptive field mapping of visual neurons (e.g. Jones & Palmer, 1987). We presented UV and green bright and dark dots of 10°visual angle at different monitor positions and recorded the corresponding population calcium activity (Fig. 3a) at the same cortical positions as shown in Fig. 2. We separated trials into quiet and active periods using the simultaneously recorded pupil size trace (quiet: <50^th^ percentile, active: >75^th^ percentile). The active periods defined by large pupil size contained all running periods (Fig. 3a). We used different pupil size thresholds for the two behavioral states compared to the model to have a sufficient number of trials in each state during the shorter recording times. For each neuron and behavioral state, we estimated a “spike”-triggered average (STA) representing the neuron’s preferred stimulus in the context of the sparse noise input (Fig. 3b; Methods). Consistent with our MEI results, most V1 neurons (81.4%) preferred a green-biased stimulus during the quiet behavioral state (Fig. 3c). Here, spectral contrast varied only slightly across the anterior-posterior axis of V1, which is likely due to the fact that we pooled data from a wider range of pupil sizes (Methods). In line with the *in-silico* analysis above, we observed a strong increase in UV-sensitivity for neurons positioned in posterior V1 during active periods (Fig. 3d and Suppl. Fig. 6e-h). This UV-shift of color preference during active periods was not observed in anterior cortex, which might be due to differences in stimulus statistics (Benjamin et al., 2019), analysis or lower signal-to-noise ratio for sparse noise responses. Together, these results confirm the CNN model’s prediction that mouse V1 color tuning changes with behavioral state, particularly for neurons sampling the upper visual field.

**Fig. 3.**
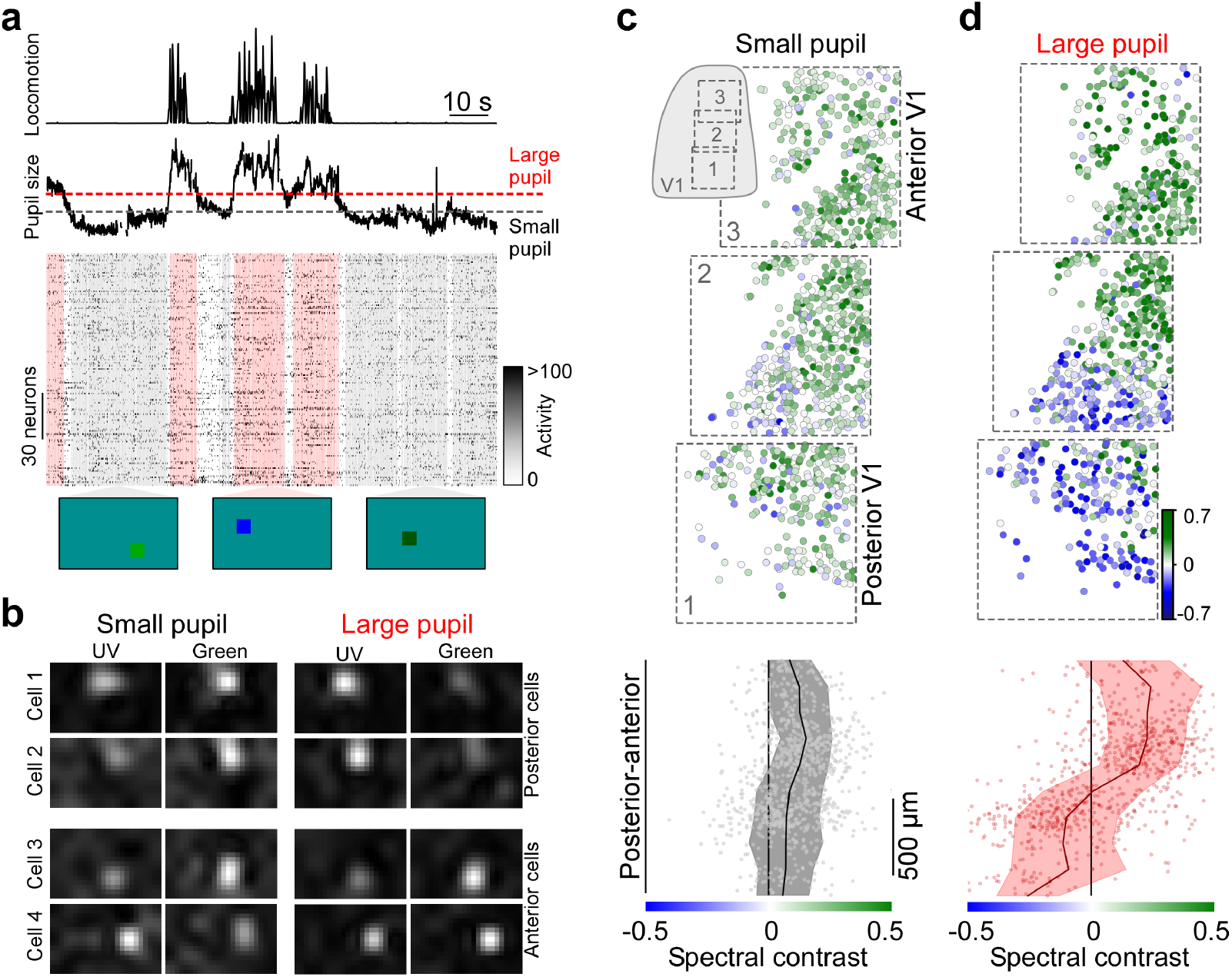
Behavioral shift of color preference for a colored sparse noise paradigm. **a**, Activity of n=150 exemplary V1 neurons in response to UV and green On and Off dots (10°visual angle) flashed for 0.2 seconds and simultaneously recorded locomotion speed and pupil size. Horizontal dotted lines indicate thresholds for small (black; <50^th^ percentile) and large (red, >75^th^ percentile) pupil trials (see Methods). Shading below highlights trials above threshold for large and small pupil. Bottom images show single stimulus frames. **b**, Spike-triggered average (STA) of 4 example neurons estimated from trials with small and large pupil, separated by posterior and anterior recording position. STAs estimated based on On and Off stimuli were combined to yield one STA per cell and pupil size (see Methods). **c**, Neurons recorded in three consecutive experiments along the posterior-anterior axis of V1 (n=981 neurons, n=3 scans, n=1 mouse), color coded based on spectral preference of their STA estimated for small pupil trials. Bottom shows spectral contrast along the posterior-anterior axis of V1 of cells from (c, top), with binned average (black, n=10 bins) and s.d. shading (gray). Spectral contrast varied significantly along anterior-posterior axis of V1 (n=981, p<0.001 for smooth term on cortical position of Generalized Additive Model (GAM); see Methods and Suppl. Statistical Analysis) **d**, Like (c), but for STAs estimated from responses to large pupil trials. Spectral contrast varied significantly along anterior-posterior axis of V1 (n=981, p<0.001 for smooth term on cortical position of GAM; see Methods and Suppl. Statistical Analysis). Optimal spectral contrast changed with pupil size (n=981, p<0.001 for behavioral state coefficient of GAM; see Methods and Suppl. Statistical Analysis), with a significant interaction between cortical position and behavioral state modulation (see Suppl. Statistical Analysis).

### Changes in pupil size mediate behavioral modulation of color tuning

Next, we investigated the mechanism underlying the observed behavior-related changes in color tuning of mouse V1 neurons. On the one hand, behavioral modulation of mouse visual responses is mediated by neuromodulatory circuits acting at different stages of the visual system (Fu et al., 2014; Lee et al., 2014; Reimer et al., 2016; Schröder et al., 2020). On the other hand, changes in pupil size during active behavioral periods result in different light intensities at the level of the retina which might also affect visual processing (Tikidji-Hamburyan et al., 2015). To experimentally test the relative contribution of these two mechanisms to the observed shift of color preference during an active state, we dissociated state-dependent neuromodulatory effects from changes in pupil size by dilating the pupil with atropine eye drops prior to recordings (Methods). During the quiet state without locomotion, this resulted in a significant increase in pupil radius (130 to 350%) compared to the control condition without pharmacological pupil dilation. We then recorded visual responses to naturalistic scenes during the dilated condition and trained a separate CNN model. The prediction performance of the model trained on data from the dilated conditions was slightly lower than the performance for control conditions (Suppl. Fig. 2a,c; 0.50 versus 0.56 correlation to average response across trials). The lower performance is likely due to the fact that, for the dilated condition, we did not incorporate pupil-related behavioral parameters and shifter network into the model due to constant pupil sizes throughout the experiment under these pharmacological conditions. We anatomically matched neurons across recording conditions based on their location after aligning each recording field to the same three-dimensional structural stack recorded after the functional imaging experiments (Methods; Walker et al., 2019a).

During the quiet state with no locomotion, the color tuning of the MEIs was systematically shifted towards higher UV-sensitivity for the recording sessions with the dilated pupil compared to the control condition (Fig. 4a-d and Suppl. Fig. 7a,b). Importantly, pharmacological pupil dilation resulted in a stronger shift of color tuning than observed during active periods under control conditions. This is likely due to the stronger pupil dilation for pharmacological (up to 15-fold increase in pupil area) compared to physiological conditions (up to 10-fold increase in pupil area). We confirmed the role of pupil size in modulating color tuning of mouse V1 neurons by recording visual responses to the sparse noise stimulus after dilating the pupil with atropine. For periods with no locomotion activity, this revealed an increased UV-sensitivity for the dilated recordings compared to control sessions (Fig. 7c-f). While the shift of color tuning with pupil dilation was stronger for posterior compared to anterior V1, there was a significant shift towards higher UV-sensitivity across all recorded V1 positions (Fig. 7f) – in line with the results predicted by the CNN model (cf. Fig. 2). Together, this suggests that pupil size changes are sufficient to shift the neurons’ color tuning towards higher UV-sensitivity.

**Fig. 4.**
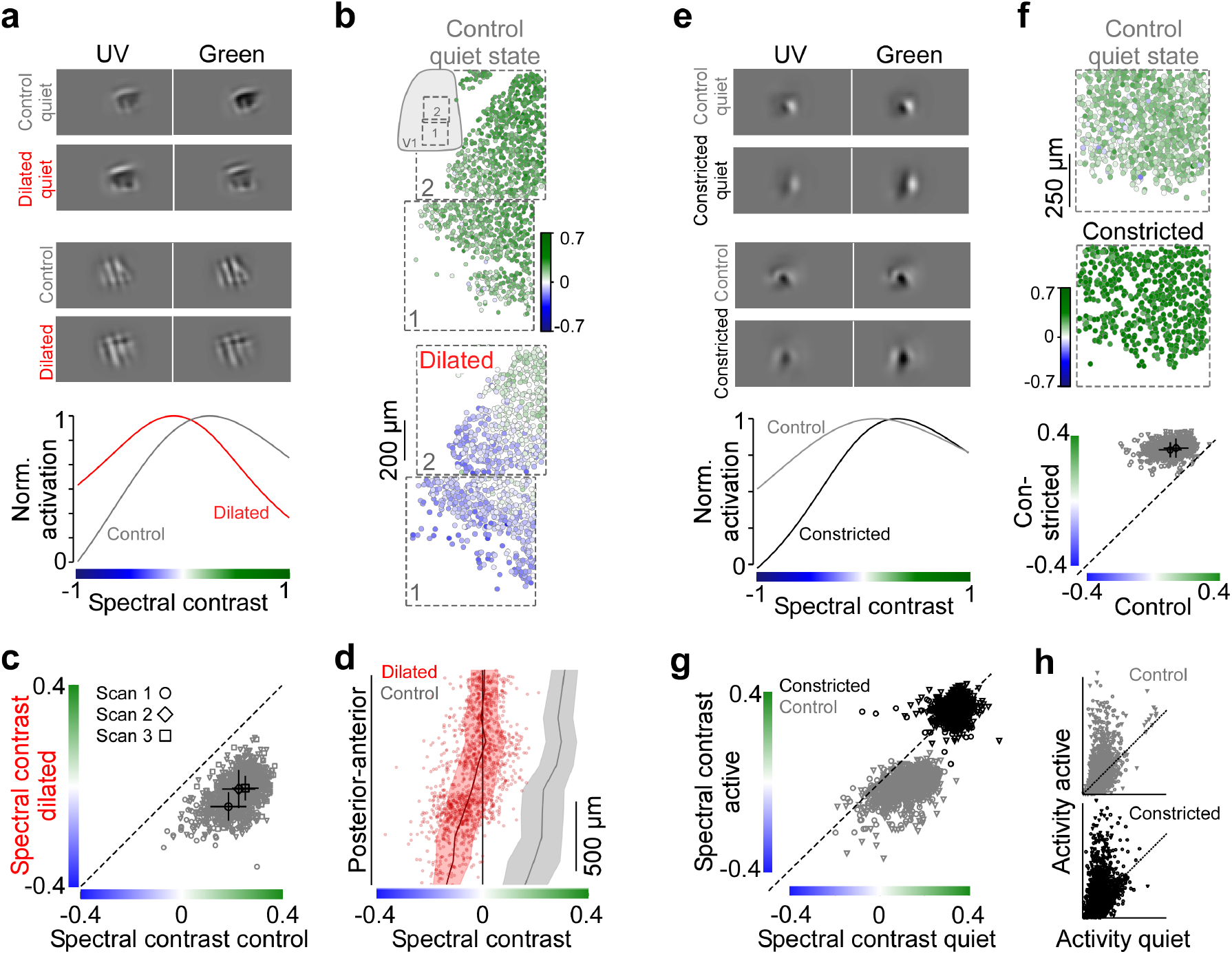
Changes in pupil size mediate shift of color tuning with behavioral state. **a**, MEIs of two example neurons optimized for a quiet state for control condition (gray) and drug condition (red) where pupil was dilated with atropine prior to recordings. Peak-normalized color tuning curves of second neuron below. Neurons were anatomically matched across recordings by alignment of the scan fields to the same 3D structural stack (see Methods). **b**, Neurons recorded in two consecutive experiments across the posterior-anterior axis of V1 for control (top; from Fig. 2) and dilated condition (bottom), color coded based on spectral contrast of MEI optimized for a quiet behavioral state. **c**, Spectral contrast of quiet state MEIs in control condition versus spectral contrast of quiet state MEIs of anatomically matched neurons in dilated condition (n=1,287 neurons, n=3 scans, n=1 mouse). Mean and s.d. of each scan indicated in black. Wilcoxon signed rank test: p<0.001 for all scans. **d**, Spectral contrast of MEIs of neurons from (b) along the posterior-anterior axis of V1 (red dots), with binned average (n=10 bins; red line) and s.d. shading. Black line and gray shading corresponds to binned average spectral contrast and s.d. of neurons recorded at the same cortical positions in control condition (cf. Fig. 2). Spectral contrast significantly varied across anterior-posterior axis of V1 for the dilated condition (n=1,859, p<0.001 for smooth term on cortical position of Generalized Additive Model (GAM); see Methods and Suppl. Statistical Analysis). Optimal spectral contrast changed with pupil dilation (n=1,859 (dilated) and n=1,759 (control), p<0.001 for condition coefficient of GAM; see Methods and Suppl. Statistical Analysis), with only a minor interaction between cortical position and behavioral state modulation (see Suppl. Statistical Analysis). **e**, Like (a), but for drug condition (black) where pupil was constricted with carbachol prior to recordings. **f**, Neurons recorded in posterior V1 (n=751 and 518 for control and constricted condition) color coded based on spectral contrast of MEI optimized for a quiet state under control (top) and constricted condition (middle). Bottom shows spectral contrast of quiet state MEIs in control versus constricted condition of anatomically matched neurons (n=1,051, n=2 scans, n=2 animals). Mean and s.d. of each scan indicated in black. Wilcoxon signed rank test: p<0.001 for all scans. **g**, Spectral contrast of MEIs optimized for a quiet state versus spectral contrast of MEIs optimized for an active state, for control (gray) and constricted condition (black). Wilcoxon signed rank test: p<0.001 for control condition, p=0.13 for constricted condition. **h**, Mean activity of n=3,379 neurons (n=2 scans, n=2 mice) during quiet and active behavioral periods in control (gray) and constricted condition (black).

Next, we studied whether the behavioral shift of color tuning is exclusively driven by pupil size. If so, constricting the pupil should (i) have an opposite effect than pupil dilation, increasing the neurons’ sensitivity to green stimuli and (ii) abolish the shift of color tuning with behavioral state. To directly test this, we recorded visual responses in posterior V1 to natural images under a control condition and while constricting the pupil with carbachol (40% of control pupil radius in the quiet state). Upon pupil constriction, pupil size did not change anymore with locomotion activity (r^2^=0.05, p=0.07; r^2^=0.52, p<0.001 for control). Similarly to the pupil dilation experiments, we trained two separate models based on control and constricted data and optimized MEIs for quiet and active behavioral states. For quiet periods, pupil constriction resulted in a systematic shift towards higher green sensitivity compared to the control condition (Fig. 4e,f). In addition, we did not observe a significant shift of color tuning during active periods for the constricted condition, while the shift was evident in the control condition (Fig. 4g). Importantly, the gain of neural activity with locomotion activity caused by neuromodulation (e.g. Fu et al., 2014) persisted in the constricted condition (Fig. 4h). Together, our experiments demonstrate that pupil size changes are both sufficient and necessary to explain V1 color tuning across behavioral states.

How might changes in pupil size increase UV-sensitivity of mouse V1 neurons? Previous studies have shown that pupil size strongly influences retinal illuminance levels in mice (Grozdanic et al., 2003; Pennesi et al., 1998), which – for specific ambient light levels – will result in different relative activation levels of rod and cone photoreceptors. Therefore, our data (cf. Figs. 2, 3) could be explained by UV- and green-sensitive cone photoreceptors taking over for active periods due to larger pupil sizes and, consequently, higher retinal light levels (Fig. 5a). To investigate this hypothesis, we computed activation levels of mouse photoreceptors as a function of pupil size based on stimulus wavelength and intensity, transmission of the mouse optical apparatus and photoreceptor sensitivity (Methods). For our experiments, we observed up to a 10-fold increase in pupil area for an active (1.9 mm^2^) compared to a quiet behavioral state (0.2 mm^2^), which resulted in a photoisomerization rate (as photoisomerizations per cone and second) of 2,000 and 200, respectively (Fig. 5b), for medium screen intensities (pixel values of 127 in 8-bit). This revealed that changes in pupil size recorded across the two behavioral states are sufficient to dynamically shift the mouse visual system from a rod- to cone-dominated operating regime (Tikidji-Hamburyan et al., 2015). Due to differences in spectral sensitivity of mouse rod and cone photoreceptors, which are especially pronounced for neurons sampling the upper visual field, this dynamic shift of activation ratios might result in an increase in UV-sensitivity. Notably, this change in spectral sensitivity caused by differential rod versus cone activation is a well-described psychophysical phenomenon in humans and other animals called Purkinje shift (Wald, 1945) – however, so far it has only been observed over much longer timescales.

**Fig. 5.**
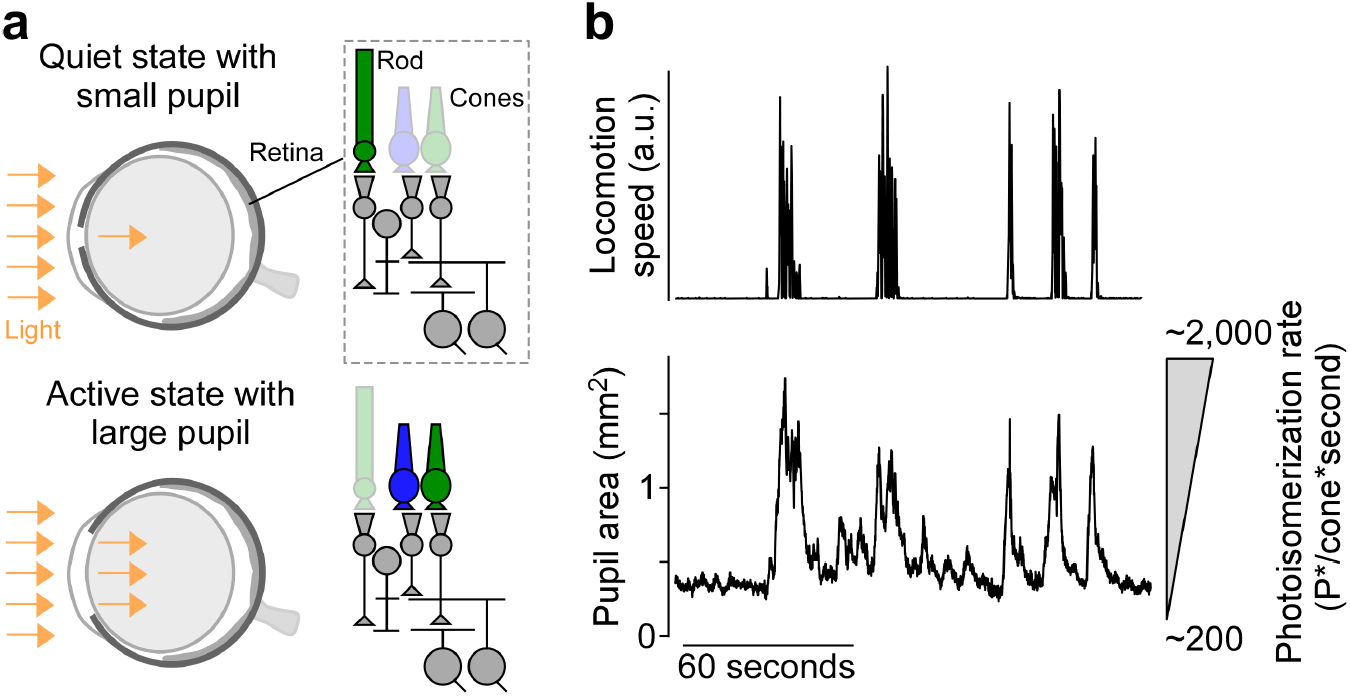
Pupil dilation during active behavioral periods greatly influences photoreceptor activation rates. **a**, Schematic of mouse eye for a quiet behavioral state with small pupil (top) and an active state with large pupil (bottom). Right panels show a simplified circuit diagram of the vertebrate retina and illustrate differential activation of rod and cone photoreceptors across the two behavioral states by varying degrees of transparency. Arrows indicate amount of light entering the eye through the pupil. Photoreceptors are color coded based on their peak wavelength sensitivity. **b**, Locomotion speed (top) and pupil area (bottom) recorded during functional imaging. Pupil dilation recorded during active behavioral periods will result in a shift from a mesopic (*≈*200 photoisomerization rate in P*/cone*second) to a low photopic regime (*≈*2,000 photoisomerization rate) for our stimulus intensities. Photoisomerization rates were estimated based on light intensity at the level of the cornea, pupil area, transmission of the optical apparatus and photoreceptor sensitivity (see Methods for detail and Franke et al., 2019; Qiu et al., 2021)

### Shift of color preference facilitates decoding of behaviorally relevant stimuli

Until now, we have characterized the behavioral modulation of mouse V1 color tuning at the single cell level and for the neurons’ optimal stimuli. Specific visual tasks, like object detection or discrimination, however, are usually solved at a population level where most neurons are not optimally stimulated. Therefore, we next tested whether the shift of color tuning during an active state might increase visual task performance at the level of an entire neuronal population and in response to natural images using an *in-silico* image reconstruction paradigm (Safarani et al., 2021). Stimulus reconstruction from neural activity has previously been used to infer the most relevant visual features encoded by the neuron population (Bialek et al., 1991; Yoshida & Ohki, 2020), like the neurons’ color sensitivity. Here we use image reconstruction with a contrast constraint on the image to analyze which dimensions in image space the neural population is more sensitive to. The underlying idea is that more important dimensions – or color channels – should be preferentially reconstructed if the reconstructed image has a limited contrast budget (see Methods and Safarani et al., 2021, for details). As the receptive fields of neurons recorded within one of our scans only covered a fraction of the screen, we used an augmented version of our CNN model for image reconstruction where the receptive field of each model neuron was copied to each pixel position of the image except the image margins (Fig. 6a; Methods). This greatly improved reconstruction performance. We presented a specific target image (image 1) to the augmented model and used the resulting response predictions to reconstruct the image (image 2) for a quiet and active state, respectively. While most reconstructed images during the quiet behavioral state exhibited higher contrast in the green compared to the UV image channel, the contrast was higher for the majority of images in the UV channel during the active state (Fig. 6b,c). This is in agreement with our results at the single cell level (cf. Figs. 2, 3) and suggests that the increase in UV-sensitivity during active periods might contribute to specific visual tasks like stimulus discrimination and detection performed by populations of neurons in mouse V1.

**Fig. 6.**
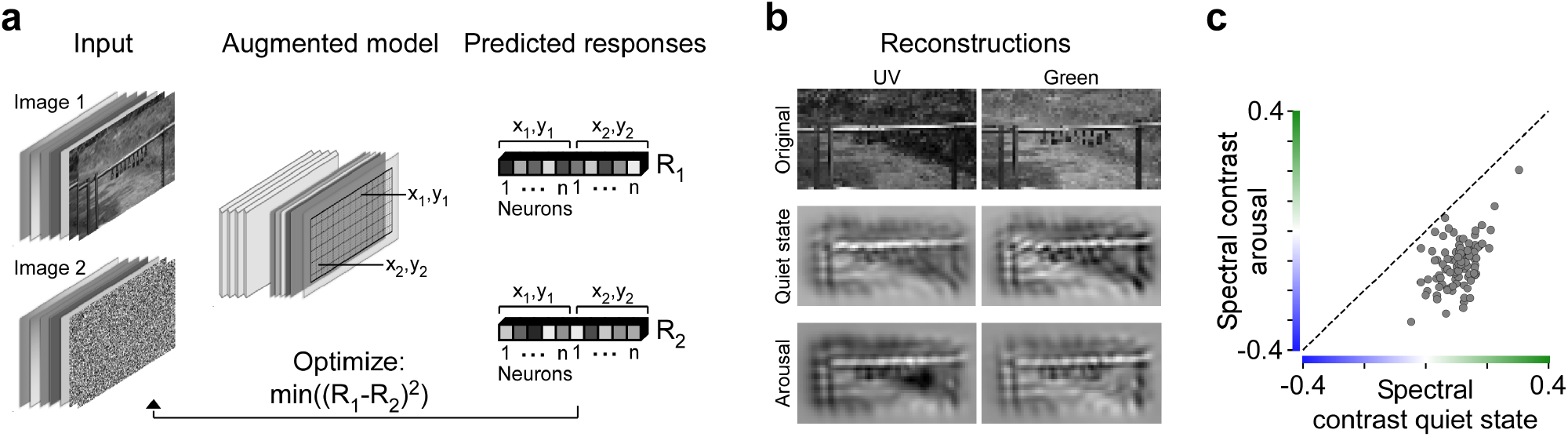
Reconstructions of colored naturalistic scenes predict color tuning shift for non-optimal stimuli of a neuron population. **a**, Schematic illustrating reconstruction paradigm. Model input is passed through an augmented model, where each model neuron is positioned at each pixel position except the image margins. For a given target input image (image 1), this results in a predicted response vector (R1) of length number of neurons times number of pixels. During image reconstruction, a novel image (image 2) is optimized such that its corresponding response vector (R2) matches the response vector of the target image as closely as possible. **b**, Green and UV image channels of exemplary test image (top) and reconstructions of this image for a quiet (middle) and active state (bottom). For reconstructions, neurons from scan 1 in Fig. 2 were used. **c**, Spectral contrasts of reconstructed test images in quiet state versus spectral contrasts of reconstructed test images in active state (n=100, p<0.001, Wilcoxon signed rank test).

We experimentally confirmed this prediction by demonstrating that neural discrimination of UV objects improves significantly during active periods. For that, we modified an object discrimination task recently described by Froudarakis et al. (2020) and presented the mouse with two different moving objects in either UV or green (Fig. 7b), while recording the population calcium activity in posterior V1 as described above. We estimated the discriminability of the two objects from the recorded neural responses using a non-linear support vector machine decoder (SVM; Methods). Specifically, we built four separate decoders with matched sample size to decode object identity during quiet and active periods from the UV and green objects, respectively (Fig. 7a), and transformed decoding accuracy into mutual information (discriminability, see Methods). Consistent with previous reports (Dadarlat & Stryker, 2017; Mineault et al., 2016; Reimer et al., 2014), we found that stimulus discriminability was higher during an active state compared to quiet behavioral periods (Fig. 7c). However, the increase in discriminability of UV objects was significantly larger than for green objects, demonstrating an increased UV sensitivity during active periods. This shift of color tuning was consistent across animals and object contrasts and polarity (Suppl. Fig. 8a-d). Only for the stimulus condition with high object contrast and no background, we found that only two of three independent recordings exhibited significantly better discriminability of UV compared to green objects during an active state (Suppl. Fig. 8d). This might correspond to a ceiling effect due to the simplicity of the task, as indicated by high object discriminability even during quiet behavioral periods.

**Fig. 7.**
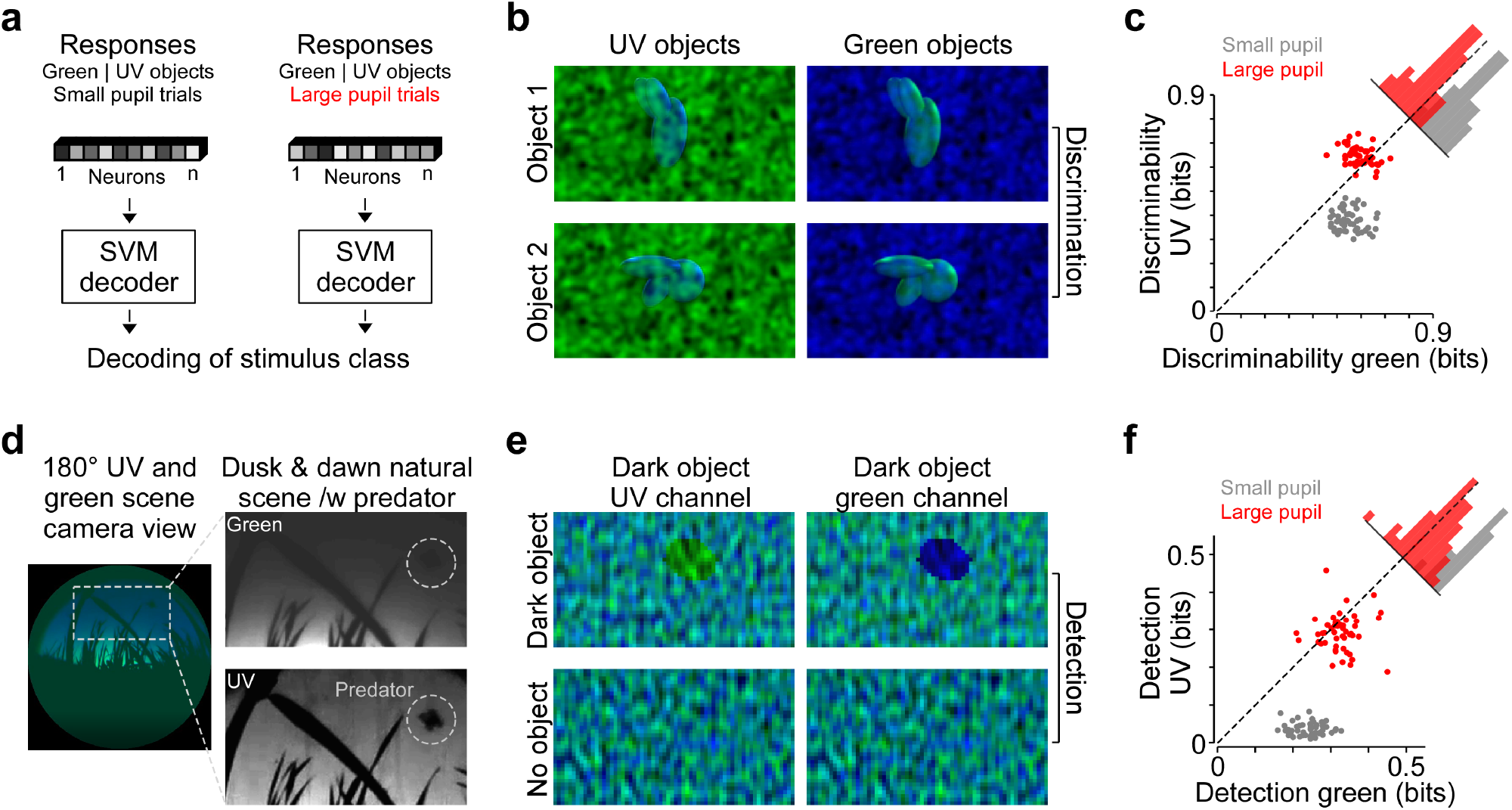
Shift of color preference facilitates decoding of behaviorally relevant stimuli. **a**, Schematic illustrating decoding paradigm. Neural responses for either quiet (small pupil) or active (large pupil) trials to green or UV objects were used to train a non-linear support vector machine (SVM) decoder to predict presented stimulus classes (see Methods for detail). **b**, Example stimulus frames of green and UV objects with UV and green noise on top, respectively, used for discrimination task. Stimulus conditions were presented as 5-second movie clips in random order (Methods). **c**, Scatter plot of discriminability of green versus UV objects for quiet (gray) and active (red) trials (Methods). Each dot represents a random sample of n=200 neurons of a neuron population (n=824) recorded in a single scan. See Suppl. Fig. 8 for more scans and stimulus conditions. Histograms represent the difference in discriminability of UV and green objects. Wilcoxon signed rank test: n=50, p<0.001. **d**, Natural scene recorded at sunrise with a custom camera adjusted to the spectral sensitivity of mice (Qiu et al., 2021), with a drone mimicking an aerial predator. Right images show single color channels of image crop from the left, with the drone highlighted by white circle. **e**, Parametric stimuli inspired by natural scene in (d), showing a dark object in either UV or green image channel. Stimulus frames with just noise (no object) were used as second stimulus condition in the detection task. Stimuli were flashed for 0.5 seconds with 0.3-0.5 second periods of gray screen in between. **f**, Similar to (c), but for detection performance of green versus UV dark objects from (d). Data shows n=50 samples of n=200 neurons from a single scan field (n=50, p<0.001, Wilcoxon signed rank test). See Suppl. Fig. 8 for an additional scan.

What might be the behavioral relevance of this increased UV sensitivity during active periods for mice? It has recently been shown that during dusk and dawn, aerial predators in the natural environment of mice are much more visible in the UV than the green wavelength range (Fig. 7d; (Qiu et al., 2021)). This is due to the fact that short wavelength UV light is more prominent in the twilight sky than green light due to ozone absorption and atmospheric scattering (discussed in Johnsen et al., 2006). Interestingly, increased scattering of UV light is also present underwater and in the snow and has been linked to specific behaviors in UV-sensitive fish (Losey et al., 1999) and arctic reindeer (Tyler et al., 2014). Therefore, an increase in UV sensitivity of mouse visual neurons for an alert behavioral state might facilitate the detection of predators visible as dark silhouettes in the sky. To test this prediction, we designed parametric stimuli inspired by these natural scenes, containing a dark ellipse of varying size, angle and position in either the green or UV image channel (Fig. 7e) on top of noise (see Methods). We adjusted the scenes with and without a dark object to have the same contrast. Then, we presented the scenes to the mouse and recorded the neural activity in posterior V1 sampling the upper visual field. Similar to the discriminability analyses above, we used four separate SVM decoders to determine detection performance for dark objects presented in the green and UV channel and during small and large pupil sizes, respectively. This revealed that detection performance of dark objects in the UV channel – the behaviorally relevant stimulus – was very low for quiet periods with small pupil, but greatly increased for an active state with larger pupil (Fig. 7f and Suppl. Fig. 8e). In contrast, detection of dark objects in the green channel did not increase during active periods to the same extent.

## Discussion

Our work identifies a novel mechanism, by which pupil dilation during active behavioral periods dynamically tunes the color sensitivity of the mouse visual system to improved detection of behaviorally relevant features. We investigated neural color tuning in mouse V1 during changes in behavioral state measured by the pupil size and locomotion combining population calcium imaging and deep neural predictive models. The resulting accurate models of neural responses allowed us to systematically characterize *in-silico* the neurons’ color tuning across different behavioral states. Our models predicted a consistent shift of color preference towards UV-sensitivity during active periods, which we confirmed experimentally. We also showed that changes in pupil size are both sufficient and necessary for the shift of color tuning during an active state, which is likely caused by a resulting switch in the engagement of rods and cones (Rhim et al., 2021). Additional decoding analyses suggested that, on the population level, the shift of feature selectivity is behaviorally relevant, as it selectively improves the detection of dark objects in the UV channel, analogous to a predatory bird flying in a UV-bright sky.

Elegant studies on invertebrates many decades ago have first demonstrated that sensory responses are modulated by the animal’s motor activity and internal state (Rowell, 1971; Wiersma & Oberjat, 1968). Since then, modulation of sensory responses as a function of behavioral and brain state has been described in many animals (e.g. Bezdudnaya et al., 2006; Maimon et al., 2010; Niell & Stryker, 2010), as well as during attention mostly studied in primates (e.g. Treue & Maunsell, 1996). State-dependent modulation predominantly affects neural responsiveness, increasing responses of some neurons while decreasing them in others (McAdams & Maunsell, 1999; Niell & Stryker, 2010; Schröder et al., 2020; Stringer et al., 2019; Zagha et al., 2013). This results in better neuronal and behavioral performance in specific visual tasks, like stimulus detection or discrimination (Bennett et al., 2013; Dadarlat & Stryker, 2017; Spitzer et al., 1988). In a few cases, however, state-dependent modulation additionally changes the tuning properties of sensory circuits. In the visual system, this has been reported, for instance, for temporal tuning in Drosophila (Chiappe et al., 2010; Jung et al., 2011), rabbits (Bezdudnaya et al., 2006), and mice (Andermann et al., 2011), as well as for direction selectivity in primates (Treue & Maunsell, 1996). In these cases, the visual system might bias processing towards visual features relevant for current behavioral goals, such as higher temporal frequencies during walking, running, and flying periods.

Here, we report a shift of neural tuning with behavioral state in mice, focusing on the color domain which has been rarely studied in the context of behavioral modulation. We demonstrate that this shift of tuning may help support ethological tasks, like the detection of predators in the sky. Consistent with this view, UV vision has been implicated in predator and prey detection in a number of animal species as an adaptation to living in different natural environments (reviewed in Cronin & Bok, 2016). This is related to stronger scattering of short wavelength light as well as ozone absorption (Hulburt, 1953) in the sky, facilitating the detection of objects as dark silhouettes against a UV bright background (Losey et al., 1999; Qiu et al., 2021). Reliable predator detection is critical for the animal’s survival at all times, including quiet states. This raises the question of how mice ensure robust predator detection during a quiet behavioral state with small pupil sizes and low UV sensitivity. A fast shift from a quiet to an active state might be initiated by brief salient stimuli, that can act as a “wake-up-call” while animals are in a quiet state (discussed in Harris & Thiele, 2011). A flying bird seen as a small moving object might correspond to such a salient stimulus.

Mechanistically, state-dependent modulation of visual responses has been linked to neuromodulators like acetlycholine and norepinephrine (reviewed in Harris & Thiele, 2011; Thiele & Bellgrove, 2018), released with active behavioral states and states of arousal. This is controlled by the basal forebrain (Berridge & Waterhouse, 2003; Foote et al., 1980; Goard & Dan, 2009), acting on neural circuits at many levels of the visual system. For example, in mouse visual cortex, release of acetylcholine via corticopetal projections activates vasoactive intestinal peptide (VIP) neurons, enhancing responses to visual stimuli (Fu et al., 2014). However, an active behavioral state, similar to a state of arousal and attention, is also associated with an increase in pupil size (Bradshaw, 1967; Hess & Polt, 1960; Reimer et al., 2014). From retinal studies, it is well established that changes in overall light level alter visual information processing (Enroth-Cugell & Lennie, 1975; Farrow et al., 2013; Tikidji-Hamburyan et al., 2015). For example, the relative strength of center and surround components of retinal receptive fields changes with the light intensity of the stimulus (Enroth-Cugell & Lennie, 1975). In principle, state-dependent changes in pupil size might therefore contribute to observed modulation of visual responses by controlling retinal illuminance levels.

Indeed, our data demonstrate that changes in pupil size are both sufficient and necessary to drive a shift of color preference of mouse V1 neurons: While pharmacological pupil dilation reproduced the spectral shift observed for physiological increases in pupil size during active periods, pupil constriction resulted in the opposite shift of color tuning. In addition, behavioral modulation of V1 color tuning was not observed while constricting the pupil, although neuromodulatory effects on response gain persisted. This strongly suggests that neuromodulation acting on the retina directly (Schröder et al., 2020) or on down-stream visual areas like dLGN (Aydın et al., 2018; Crombie et al., 2021) and V1 (Niell & Stryker, 2010; Reimer et al., 2014; Vinck et al., 2015) is not necessary to evoke the shift of color tuning during active periods. We propose that larger pupil sizes during active periods increase the amount of light entering the eye to an extent that can initiate a shift from rod- to cone-dominated vision, at least for mesopic ambient light levels. Earlier studies have measured the range of pupil sizes in mice (Grozdanic et al., 2003; Pennesi et al., 1998) and found that an increase from smallest (0.1mm^2^) to largest (4.3 mm^2^) pupil area will alter retinal illuminance levels by ≈1.5 orders of magnitude. A recent neurophysiological study on anaesthetized mice demonstrates that pharmacological pupil dilation at constant ambient light levels is sufficient to induce a shift from rod- to cone-driven visual responses in V1 (Rhim et al., 2021). Our data indicates that a switch between the rod and cone system can also happen dynamically in behaving mice as a consequence of changes in pupil size across distinct behavioral states. Cone activation rates measured in mouse (Joachimsthaler & Kremers, 2019) and primate retina (van Hateren & Lamb, 2006) support a fast switch from rod- to cone-driven visual responses for timescales below one second. As rod and cone photoreceptors differ with respect to light sensitivity, temporal resolution, and degree of non-linearity (discussed in Lamb, 2016), dynamically adjusting their relative activation will greatly influence the sensory representation of the visual scene. Together, this suggests that in addition to neuromodulators and other state-dependent top-down effects in cortex, dynamic changes in pupil size can affect cortical tuning through a mechanism acting already at the very first stage of the visual system – the rod and cone photoreceptors.

Our findings provide new insights to the long standing debate of the functional role of pupil dilation associated with states like arousal and attention present across many animal species (reviewed in Joshi & Gold, 2020; Larsen & Waters, 2018). It has been speculated that higher amounts of light entering the eye might improve visual performance. At the same time, the pupil size for a distinct ambient light level is optimized for both sensitivity and visual acuity (Laughlin, 1992). Therefore, an additional increase in pupil size might decrease visual acuity. However, this is only relevant for species like primates where visual acuity is limited by optical aberrations rather than neurophysiological parameters (Green, 1970; Williams & Coletta, 1987), like photoreceptor density and retinal convergence. Our results propose a new role for state-dependent pupil dilation: To tune the visual system to specific behavioral tasks by dynamically adjusting the relative activation of rod and cone photoreceptors, as demonstrated here for predator detection in mice during dusk and dawn. Next to ambient light levels, the effect of pupil size on the differential activation of the cone and rod system will depend on how much the pupil diameter can influence retinal illuminance levels, i.e. the size of the pupil relative to the size of the eye, as well as the contribution of behavioral state in modulating pupil diameter. For most animal species, this information is not yet available. Interestingly, pupil dilation is likely under voluntary control for some animals such as birds and reptiles (discussed in Douglas, 2018), and potentially even for some humans (Eberhardt et al., 2021). Together, this opens up many exciting questions about how changes in pupil size contribute to visual information processing across animal species.

## ACKNOWLEDGEMENTS

We thank Greg Horwitz, Thomas Euler, Mackenzie Mathis, Tom Baden, Lara Höfling and Yongrong Qiu for feedback on the manuscript and Donnie Kim, Daniel Sitonic, Dat Tran, Zhuokun Ding, Konstantin Lurz, Mohammad Bashiri, Christoph Blessing and Edgar Walker for technical support and discussions. This work was supported by the Carl-Zeiss-Stiftung (FS), the DFG Cluster of Excellence “Machine Learning – New Perspectives for Science” (FS; EXC 2064/1, project number 390727645), the AWS Machine Learning research award (FS), and the Intelligence Advanced Research Projects Activity (IARPA) via Department of Interior/Interior Business Center (DoI/IBC) contract number D16PC00003 (AST). The U.S. Government is authorized to reproduce and distribute reprints for Governmental purposes notwithstanding any copyright annotation thereon. Disclaimer: The views and conclusions contained herein are those of the authors and should not be interpreted as necessarily representing the official policies or endorsements, either expressed or implied, of IARPA, DoI/IBC, or the U.S. Government. Also supported by R01 EY026927 (AST), NEI/NIH Core Grant for Vision Research (T32-EY-002520-37) and NSF NeuroNex grant 1707400 (AST).

## AUTHOR CONTRIBUTIONS

**KF**: Conceptualization, Methodology, Validation, Software, Formal Analysis, Investigation, Writing - Original Draft, Visualization, Supervision, Project administration; **KW**: Conceptualization, Methodology, Validation, Software, Formal Analysis, Investigation, Writing - Original Draft, Visualization, Data Curation; **KP**: Investigation, Validation, Writing - Review & Editing; **MG**: Investigation, Validation; **TM**: Investigation; **SP**: Methodology, Software, Validation, Writing - Review & Editing; **EF**: Methodology, Investigation, Writing - Review & Editing; **JR**: Validation, Writing - Review & Editing; **FS**: Conceptualization, Writing - Review & Editing, Methodology, Software, Data curation, Supervision, Funding acquisition; **AT**: Conceptualization, Experimental and analysis design, Supervision, Funding acquisition, Writing - Review & Editing

## Materials and Methods

### Neurophysiological experiments

All procedures were approved by the Institutional Animal Care and Use Committee of Baylor College of Medicine. Mice of either sex (Mus musculus, n=6) expressing GCaMP6s in excitatory neurons via Slc17a7-Cre and Ai162 transgenic lines (stock number 023527 and 031562, respectively; The Jackson Laboratory) were anesthetized and a 4 mm craniotomy was made over the visual cortex of the right hemisphere as described previously (Froudarakis et al., 2014; Reimer et al., 2014). For functional recordings, awake mice were headmounted above a cylindrical treadmill and calcium imaging was performed using a Ti-Sapphire laser tuned to 920 nm and a two-photon microscope equipped with resonant scanners (Thorlabs) and a 25x objective (MRD77220, Nikon). Laser power after the objective was kept below 60mW. The rostro-caudal treadmill movement was measured using a rotary optical encoder with a resolution of 8,000 pulses per revolution. We used light diffusing from the laser through the pupil to capture eye movements and pupil size. Images of the pupil were reflected through a hot mirror and captured with a GigE CMOS camera (Genie Nano C1920M; Teledyne Dalsa) at 20 fps at a 1,920 × 1,200 pixel resolution. The contour of the pupil for each frame was extracted using DeepLabCut (Mathis et al., 2018) and the center and major radius of a fitted ellipse were used as the position and dilation of the pupil.

To identify V1 boundaries, we used pixelwise responses to drifting bar stimuli of a 2,400 x 2,400 μm scan at 200 μm depth from cortical surface (Garrett et al., 2014), recorded using a large field of view mesoscope (Sofroniew et al., 2016) not used for other functional recordings. Imaging was performed in V1 using 512 x 512 pixel scans (650 x 650 μm) recorded at approx. 15 Hz and positioned within L2/3 at around 200 μm from the surface of the cortex. Imaging data were motion-corrected, automatically segmented and deconvolved using the CNMF algorithm (Pnevmatikakis et al., 2016); cells were further selected by a classifier trained to detect somata based on the segmented cell masks. In addition, we excluded cells with low stimulus correlation. For that, we computed the first principal component (PC) of the response matrix of size number of neurons x number of trials and then for each neuron estimated the linear correlation of its responses to the first PC, as the first PC captured unrelated background activity. We excluded neurons with a correlation lower or higher than −0.25 or 0.25, respectively. This resulted in 450–1,100 selected soma masks per scan depending on response quality and blood vessel pattern. A structural stack encompassing the scan plane and imaged at 1.6 × 1.6 × 1 μm xyz resolution with 20 repeats per plane was used to register functional scans into a shared xyz frame of reference. Cells registered to the same three-dimensional stack were then matched for distances of <10 μm. For inception loop experiments, we confirmed the anatomical matching with a functional matching procedure, using the cells’ responses to the same set of test images (see also (Walker et al., 2019a)) and only included matched neurons with a response correlation of >0.5 for further analysis. To bring different recordings across the posterior-anterior axis of V1 into the same frame of reference, we manually aligned the mean image of each functional recording to the mean image of the 2,400 x 2,400 μm scan acquired at the mesoscope (see above) using the blood vessel pattern. Then, each cell within the functional scan was assigned a new xy coordinate (in μm) in the common frame of reference.

### Visual stimulation

Visual stimuli were presented to the left eye of the mouse on a 42 x 26 cm light-transmitting teflon screen (McMaster-Carr) positioned 12 cm from the animal. Light was back-projected onto the screen by a DLP-based projector (EKB Technologies Ltd; Franke et al., 2019) with UV (395 nm) and green (460 nm) LEDs that differentially activated mouse S- and M-opsin. LEDs were synchronized with the microscope’s scan retrace. Light intensity (as photoisomerization rate, P* per second per cone) was calibrated using a spectrometer (USB2000+, Ocean Optics) to result in equal activation rates for mouse M- and S-opsin (for details, see (Franke et al., 2019; Qiu et al., 2021) and calibration notebook). In brief, spectrometer readings were transformed into photon flux and then photoisomerization rate, considering both the wavelength-specific transmission of the mouse optical apparatus (Henriksson et al., 2010) and the ratio between pupil size and retinal area (Schmucker & Schaeffel, 2004). For a pupil area of 0.2 mm^2^ during quiet trials and maximal stimulus intensities (255 pixel values), this resulted in 400 P* per second and cone type corresponding to the mesopic range. During active periods, the pupil area increased to 1.9 mm^2^ resulting in 4,000 P* per second and cone type corresponding to the low photopic regime.

Prior to functional recordings, the screen was positioned such that the population receptive field across all neurons within a 50 x 50 μm central patch, estimated using an achromatic sparse noise paradigm, was within the center of the screen. Screen position was fixed and kept constant across recordings of the same neurons. We used Psychtoolbox in Matlab for stimulus presentation and showed the following light stimuli:

#### Natural images

We presented naturalistic scenes from an available online database (Russakovsky et al., 2015). We selected images based on two criteria (see also Suppl. Fig. 1). First, to avoid an intensity bias in the stimulus, we selected images with no significant difference in mean intensity of blue and green image channel across all images. Second, we selected images with high pixelwise mean squared error (MSE >85) across color channels to increase chromatic contrast, resulting in lower pixel-wise correlation across color channels compared to a random selection. Then, we presented the blue and green image channel using the UV and green LED of the projector, respectively. For a single scan, we presented 4,500 unique colored and 750 monochromatic images in UV and green, respectively. We added monochromatic images to the stimulus to include images without correlations across color channels, thereby diversifying the input to the model. As test set, we used 100 colored and 2 x 25 monochromatic images that were repeated 10 times uniformly spread throughout the recording. Each image was presented for 500 ms, followed by a gray screen (UV and green LED at 127 pixel value) for 300 to 500 ms, sampled uniformly from that range.

#### Sparse noise

To map the receptive fields of V1 neurons, we used a sparse noise paradigm. UV and green bright (pixel value 255) and dark (pixel value 0) dots of approx. 10° visual angle were presented on a gray background (pixel value 127) in randomized order. Dots were presented for 8 and 5 positions along the horizontal and vertical axis of the screen, respectively, excluding screen margins. Each presentation lasted 200 ms and each condition (e.g. UV bright dot at position x=1 and y=1) was repeated 50 times.

#### Full-field binary white noise

We used a binary full-field binary white noise stimulus of UV and green LED to estimate temporal kernels of V1 neurons. For that, the intensity of UV and green LED was determined independently by a balanced 15-minute random sequence updated at 10 Hz. A similar stimulus was recently used in recordings of mouse (Szatko et al., 2020) and zebrafish retina (Yoshimatsu et al., 2020).

#### Colored objects

To test for object discrimination, we used two synthesized objects rendered in Blender (www.blender.org) as described recently (Froudarakis et al., 2020). In brief, we smoothly varied object position, size, tilt, and axial rotation. For bright objects, we also varied either the location or energy of 4 light sources. Stimuli were either rendered as bright objects on a black screen and Gaussian noise in the other color channel (condition 1), bright and dark objects on a gray screen and Gaussian noise in the other color channel (condition 2 and 3) or as bright objects on a black screen without Gaussian noise (condition 4). Per object and condition, we rendered movies of 875 seconds, which we then divided into 175 5-second clips. We presented the clips with different conditions and objects in random order.

#### Images with dark objects

For the object detection task, we generated images with independent Perlin noise (Perlin, 1985) in each color channel using the perlin-noise package for Python. Then, for all images except the noise images, we added a dark ellipse (pixel value 0) of varying size, position, and angle to one of the color channels. We adjusted the contrast of all images with a dark object to match the contrast of noise images, such that the distribution of image contrasts did not differ between noise and object images. We presented 2,000 unique noise images and 2,000 unique images with a dark object in the UV and green image channel, respectively. Each image was presented for 500 ms, followed by a gray screen (UV and green LED at 127 pixel value) for 300 to 500 ms, sampled uniformly from that range.

For the presentation of naturalistic scenes and object movies and images, we applied a gamma function of 1.9 to the 8-bit pixel values of the monitor.

### Preprocessing of neural responses and behavioral data

Neural responses were first deconvolved using constrained non-negative calcium deconvolution (Pnevmatikakis et al., 2016). For all stimulus paradigms except the full-field binary white noise stimulus, we subsequently extracted the accumulated activity of each neuron between 50 ms after stimulus onset and offset using a Hamming window. For the presentation of objects, we segmented the 5-second clips into 9 bins of 500 ms, starting 250 ms after stimulus onset. Behavioral traces were extracted using the same temporal offset and integration window as deconvolved calcium traces. To train our models, we isotropically downsampled stimuli images to 64 × 36 pixels. Input images, the target neuronal activities, behavioral traces and pupil positions were normalized across the training set during training.

### Pharmacological manipulations

To dilate and constrict the pupil pharmacologically, we applied 1-3% atropine and carbachol eye drops, respectively, to the left eye of the animal facing the screen for visual stimulation. Functional recordings started after the pupil was dilated or constricted. Pharmacological pupil dilation lasted >2 hours, allowing to use all data for further analysis. In contrast, carbachol eye drops constricted the pupil for approx. 30 minutes and were re-applied once during the scan. For analysis, we then only selected trials with constricted pupil (n=1.800 and n=2.400 for animal 1 and 2, respectively) and matched data analyzed in the control scans to the same trial numbers.

### Sparse noise spatial receptive field mapping

We estimated spatial spike-triggered averages (STAs) of V1 neurons in response to the sparse noise stimulus by multiplying the stimulus matrix with the response matrix of each neuron (Schwartz et al., 2006), separately for each stimulus color and polarity as well as behavioral state. For the latter, we separated trials into small (< 50th percentile) and large pupil trials (> 75th percentile). We used different pupil size thresholds for the two behavioral states compared to the model due to shorter recording time. For recordings with pupil dilation, we used locomotion speed instead of pupil size to separate trials into two behavioral states. For each behavioral state, STAs computed based on On and Off dots were averaged to yield one STA per cell and stimulus color. Green and UV STAs of the same behavioral state were peak-normalized to the same maximum. To assess STA quality, we generated response predictions by multiplying the flattened STA of each neuron with the flattened stimulus frames and compared the predictions to the recorded responses by estimating the linear correlation coefficient. For analysis, we only included cells where correlation >0.2 for at least one of the stimulus conditions. In contrast to the modelling results, STA spectral contrast for a quiet state did only slightly vary across anterior-posterior axis of V1. We verified that this is due different pupil size threshold by using the data in response to natural images (cf. Fig. 2) to train a separate model without behavior as input channels on trials with small pupil (<50^th^ percentile) and subsequently optimized MEIs – a procedure more similar to the STA paradigm. When looking at the spectral contrast of the resulting MEIs, we indeed observed a smaller variation of color preference across the anterior-posterior axis of V1, confirming our prediction.

### Full-field binary white noise temporal receptive field mapping

We used the responses to the 10 Hz full-field binary white noise stimulus of UV and green LED to compute temporal STAs of V1 neurons. Specifically, we upsampled both stimulus and responses to 60 Hz and then multiplied the stimulus matrix with the response matrix of each neuron. Per cell, this resulted in a temporal STA in response to UV and green flicker, respectively. Kernel quality was measured by comparing the variance of each temporal STA with the variance of the baseline, defined as the first 100 ms of the STA. Only cells with at least 5-times more variance of the kernel compared to baseline were considered for further analysis.

### Simulated data using Gabor neurons

We simulated neurons with Gabor receptive fields with varying Gabor parameters across the two color channels. Then, we normalized each Gabor receptive field to have a background of 0 and an amplitude range between −1 and 1. To generate responses of simulated neurons, we used the same set of training images presented during functional recordings. First, we subtracted the mean across all images from the training set, multiplied each Gabor receptive fields with each training image and computed the sum of each multiplication across the two color channels *c*. We then passed the resulting scalar response per neuron through a rectified linear unit (ReLU), to obtain the simulated response *r*, such that:

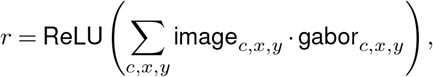

where

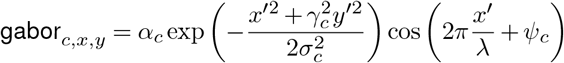

with *x*′ = *x* cos(*θ_c_*) + *y* sin(*θ_c_*) and *y*′ = −*x* sin(*θ_c_*) + *y* cos(*θ_c_*). We varied orientation *θ*, size *σ*, spatial aspect ratio *γ*, phase *Ψ*, and color preference *α* indepentenly for each color channel and neuron, while keeping spatial frequency *λ* constant across all neurons. Finally, we passed the simulated responses *r* through a Poisson process and normalized the responses by the respective standard deviation of the responses across all images. We used the responses of the simulated Gabor neurons together with the natural images to train the model (see below). Our model recovered both color-opponency and color preference of simulated neurons. Only extreme color preferences were slightly underestimated by our model, which is likely due to high correlations across color channels of natural scenes.

### Network architecture

To perform our in *in-silico* tuning characterization, we created a convolutional neural network (CNN) model, which was split into two parts: the *core* and the *readout*. The core computed latent features from the inputs, which were shared among all neurons. The readout was learned per neuron and mapped the output features of the core onto the neuronal responses via regularized regression.

#### Representation/Core

We based our model on the work of (Lurz et al., 2021), as it was demonstrated to set the state of the art for predicting the responses of a population of mouse V1 neurons. In brief, we modelled the core as a four-layer CNN, with 64 feature channels per layer. Each layer consisted of a 2d-convolutional layer followed by a batch normalization layer and an ELU nonlinearity (Clevert et al., 2015; Ioffe & Szegedy, 2015). Except for the first layer, all convolutional layers were depth-separable convolutions (Chollet, 2017) which lead to better performance while reducing the number of core parameters. Each depth-separable layer consisted of a 1×1 point-wise convolution, followed by a 7×7 depthwise convolution, again followed by a 1×1 pointwise convolution. Without stacking the outputs of the core, the output tensor of the last layer was passed on to the readout.

#### Readouts

To get the scalar neuronal firing rate for each neuron, we computed a linear regression between the core output tensor of dimensons **x** ∈ ℝ^*w×h×c*^ (**w**idth, **h**eight, **c**hannels) and the linear weight tensor **w** ∈ ℝ^*c×w×h*^, followed by an ELU offset by one (ELU+1), to keep the response positive. We made use of the recently proposed Gaussian readout (Lurz et al., 2021), which simplifies the regression problem considerably. Our Gaussian readout learned the parameters of a 2D Gaussian distribution 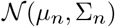 and sampled a location of height and width in the core output tensor in each training step, for every image and neuron. Given a large enough initial Σ_*n*_ to ensure gradient flow, Σ_*n*_, i.e. the uncertainty about the readout location, decreased during training for more reliable estimates of the mean location *μ_n_*, which represented the center of a neuron’s receptive field. At inference time (i.e. when evaluating our model), we set the readout to be deterministic and to use the fixed position *μ_n_*. We thus learned a position of a single point in core feature space for each neuron. In parallel to learning the position, we learned the weights of the weight tensor of the linear regression of size *c* per neuron. Furthermore, we made use of the retinotopic organization of V1, by coupling the recorded cortical 2d-coordinates **p**_*n*_ ∈ ℝ^2^ of each neuron with the estimation of the receptive field position *μ_n_* of the readout. We achieved this by learning a common function *μ_n_* = *f* (**p**_*n*_), which is shared by all neurons. We set *f* to be a randomly initialized linear fully connected network of size 2-2 followed by a *tanh* nonlineartiy.

#### Shifter network

Because we used a free viewing paradigm when presenting the visual stimuli to the head-fixed mice, the RF positions of the neurons with respect to the presented images had considerable trial to trial variability. To inform our model of the trial dependent shift of the neurons receptive fields, we shifted *μ_n_*, the model neuron’s receptive field center, using the estimated pupil center (see section Neurophysiological experiments above). We accomplished this by passing the pupil center through a small shifter network, a three layer fully connected network with *n* = 5 hidden features, again followed by a *tanh* nonlineartiy, that calculates the shift Δx and Δy per trial. The shift is then added to *μ_n_* of each model neuron.

#### Input of behavior and image position encoding

In addition to the green and UV channel of the visual stimulus, we appended five extra channels to each input to the model. We added three channels of the recorded behavioral parameters in each given trial (pupil size, instantaneous change of pupil size, and running speed), such that each channel simply consisted of the scalar for the respective behavioral parameter, transformed into stimulus dimension. This enabled the model to predict neural responses as a function of both visual input and behavioral parameters in each given trial (pupil size, instantaneous change of pupil size, and running speed), such that each channel simply consisted of the scalar for the respective behavioral parameter, transformed into stimulus dimension. This enabled the model to predict neural responses as a function of both visual input and behavior and thus to learn the relationship between behavioral states and neuronal activity. This modification allowed us to investigate the effect of behavior by selecting different inputs in the behavioral channels while keeping the image unchanged. Furthermore, we added a positional encoding to the inputs, consisting of two channels, encoding the horizontal and vertical pixel positions of the visual stimulus. These encodings can be thought of as simple grayscale gradients in either direction, with values from [−1, …, 1]. Appending position encodings of this kind have been shown to improve the ability of CNNs to learn spatial relationships between pixel positions of the input image and high level feature representations (Liu et al., 2018). We found that including the position embedding increased the performance of our model (Suppl. Fig. 2b). We also observed a smoother gradient of color tuning across the different scan fields (2b, c) when adding the position encoding, indicating that indeed the model learned the color sensitivity tuning of mouse V1 more readily.

### Model training and evaluation

We first split the unique training images into the training and validation set, using a split of 90% to 10%, respectively. Then we trained our networks with the training set by minimizing the Poisson loss 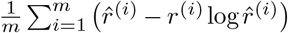, where *m* denotes the number of neurons, 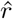 the predicted neuronal response and *r* the observed response. After each full pass through the training set (i.e. epoch), we calculated the correlation between predicted and measured neuronal responses across all neurons on the validation set: if the correlation failed to increase during a fixed number of epochs, we stopped the training and restored the model to its state after the best performing epoch. After each stopping, we either decreased the learning rate or stopped training altogether, if the number of learning-rate decay steps was reached. We optimized the network’s parameters via stochastic gradient descent using the Adam optimizer (Kingma & Ba, 2015). Furthermore, we performed an exhaustive hyper-parameter selection using Bayesian search on a held-out dataset. All parameters and hyper-parameters can be found in our GitHub repository (see Code Availability). When evaluating our models on the test set (Suppl. Fig. 2), we used two different types of correlation. Firstly, referred to as test correlation, we computed the correlation between the model’s prediction and neuronal responses across single trials, including the trial by trial variability across repeats. Secondly, we computed the correlation of the predicted responses with the average responses across repeats and refer to it here as correlation to average. We also computed the fraction of variance explained, using 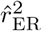 proposed by (Pospisil & Bair, 2020), which provides an unbiased estimate of the variance explained based on the expected neuronal response across image repetitions. However, our model computed different predictions for each repetition of a given test set image, because we also fed the behavioral parameters of each trial into the model. We thus simply averaged the model responses across repetitions and calculated the 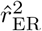 accordingly.

### Ensemble models

For all analyses as well as for the generation of MEIs, we used an ensemble of models, rather than individual models. For each dataset, we trained 10 individual models that were initialized with different random seeds. We then selected the best 5 models as measured by their performance on the validation set to be part of the ensemble. The inputs to the ensemble model were passed to each member, and the resulting predictions were averaged to obtain the final model prediction.

### Generation of maximally exciting inputs

We used a variant of regularized gradient ascent on our trained deep neural network models to obtain a maximally exciting input image (MEI) for each neuron, given by **x** ∈ ℝ^*h×w×c*^, with height *h*, width *w*, and channels *c*. Because of our particular model inputs (see section Input of behavioral parameters and image position encoding), each MEI, like the natural images used for training, had seven channels, of which we optimize only the first two: the green and UV color channels. To obtain MEIs, we initialized a starting image with Gaussian white noise. We set the behavioral channels of the starting image to the desired behavioral values, as well as setting the position channels to the default position encoding. Then, in each iteration of our gradient ascent method, we showed the image to the model and computed the gradients of the first two image channels (green and UV) w.r.t. the model activation of a single neuron. During gradient descent optimization, we smoothed the gradient by applying Gaussian blur with a *σ* of 1 pixel. To constrain the contrast of the image, we calculated the Euclidean (L2) norm of the resulting MEI

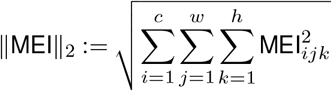

across all pixels MEI_*ijk*_ of the two color channels *c* and compared the L2 norm to a fixed norm budget *b*, which we set to 10. The norm budget can be effectively thought of as a contrast constraint. An L2 norm of 10, calculated across all pixel intensities of the image, proved to be optimal such that the resulting MEI had minimal and maximal values similar to those found in our training natural image distribution. If the image exceeded the norm budget during optimization, we divided the entire image by factor *f_norm_* with *f_norm_* = ∥MEI∥_2_/*b*. Additionally, we made sure that the MEI could not contain values outside of the 8-bit pixel range by clipping the MEI outside of these bounds, corresponding to 0 or 255 pixel-intensity. As an optimizer, we used stochastic gradient descent with learning rate of 3. We ran each optimization for 1000 iterations, without an option for early stopping. Our analyses showed that the resulting MEIs were highly correlated across behavioral states (Suppl. Fig. 5a-c). To validate this finding, we performed a control experiment using two separate models exclusively trained on trials from active or quiet states. We again split the trials into quiet and active periods using pupil size (quiet: <50^th^ percentile, active: >75^th^ percentile). When inspecting the MEIs generated from these two models, we found that the MEIs were again highly correlated across color channels, albeit less than for the model that was trained on the entire data. This can partly be explained by the limited amount data for the model trained with trials from the active state that occurred less frequently in our data.

### Spectral contrast

For estimating the chromatic preference of the recorded neurons, we used spectral contrast (*SC*). It is estimated as Michelson contrast ranging from −1 and 1 for a neuron responding solely to UV and image, respectively. We define *SC* as

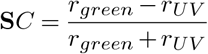

where *r_green_* and *r_UV_* correspond to (i) the norm of green and UV MEI channel to estimate the neurons’ chromatic preference in the context of naturalistic scenes, (ii) the amplitude (mean of all pixels >90^th^ percentile) of UV and green spatial STA to estimate the neurons’ chromatic preference in the context of the sparse noise paradigm, (iii) the norm of the green and UV channel of reconstructed images to quantify chromatic preference at a populational level and (iv) the norm of green and UV channel of simulated Gabor RFs to obtain each simulated neuron’s chromatic preference.

### In silico color tuning curves

To generate in silico color tuning curves for recorded V1 neurons, we systematically varied the L2-norm of the green and UV MEI channel while keeping the overall norm across color channels constant (with norm = 10). We used n=50 spectral contrast levels, ranging from all contrast in the UV channel to all contrast in the green channel. We then showed the modified MEIs to the model and plotted the predicted responses across all n=50 spectral contrast levels. Modified MEIs were either presented to the model for a quiet or active state (see also above).

### Reconstruction analysis

We visualize which image features the population of model neurons are sensitive to by using a novel resource constrained image reconstruction method based on the responses of a population of model neurons. (Safarani et al., 2021). The reasoning behind a resource constrained reconstruction is to recreate the responses of a population of neurons when presented with a target image, by optimizing a novel image and matching the neuron’s responses given that novel image as close as possible to the target image’s responses. By limiting the image contrast of the reconstructed image during the optimization, the reconstructions will only contain the image features that are most relevant to recreate the population responses, thereby visualizing the sensitivities and invariances of the population of neurons. As target images for our reconstruction, we chose natural images from our test set. For each reconstruction, we first calculated the neuronal responses *f* (**x**_0_) of all model neurons when presented with target image **x**_0_. We then initialized an image (**x**) with Gaussian white noise as the basis for reconstruction of the target image by minimizing the squared loss between the target responses and the responses from the reconstructed image *ℓ* (**x**_0_, **x**) = ∥*f* (**x**) − *f* (**x**_0_)∥^2^ subject to a norm constraint. In this work, we set the contrast (i.e. L2-norm, see section Generation of maximally exciting inputs for details) of the reconstructions to 40, which corresponds to ~60% of the average norm of our natural image stimuli. We chose this value to be high enough to still allow for qualitative resemblance between the the reconstructed image and the target, while keeping the constraint tight enough to avoid an uninformative trivial solution, i.e. the identical reconstruction of the target. We improved the quality of the reconstructions by using an augmented version of our model, which reads out each neuronal response not from the model neuron’s actual receptive field position *μ* (see Readouts for details), but instead from all height-times-width positions in feature space, except the n=10 pixels around each border to avoid padding artefacts. This yielded 18 ∗ 46 = 828 copies per neuron and with the N=478 original model neurons, this resulted in overall n=395,784 augmented neurons for our reconstruction analyses. We found stochastic gradient descent with a learning rate of 1000 to yield the qualitatively best reconstructions, resulting in images with the least amount of noise. We always optimized for 5000 steps per image, without the early stopping the optimization.

### Decoding analysis

We used a support vector machine classifier with a radial basis function kernel to estimate decoding accuracy between the neural representations of two stimulus classes - either object 1 and object 2 (object discrimination) or dark object and no object (object detection). For that, we randomly sampled n=200 neurons from all neurons recorded within one scan. For a given neuron selection, we build four separate decoders for UV and green stimuli and small and large pupil trials, respectively. Then, we train each decoder with n=175 randomly selected training trials, test its accuracy using n=25 randomly selected test trials and compute the mean accuracy across n=10 different training/test trial splits. We repeat this procedure for n=50 different neuron selections. Finally, we convert the decoding accuracy into discriminability, the mutual information between the true class and its estimate using

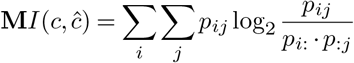

where *p_ij_* is the probability of observing the true class *i* and predicted class *j* and *p*_*i*:_ and *p*_:*j*_ denote the respective marginal probabilities.

### Response reliability

We calculated the signal-to-noise ratio (SNR) as our measure for response reliability. Following (Pospisil & Bair, 2020), we defined the SNR as follows:

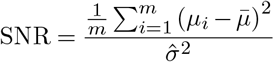

This measure of a neuronal turning curve expresses the ratio of the variance in the expected responses against trial-by-trial variability across repeats, with *μ_i_* being the expected response to the *i*th stimulus, and the average expected response

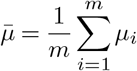

The trial-by-trial variance 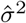 is given as the average variance across repeats over all stimuli. Importantly, we assume that 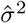 is constant across all responses to different stimuli. This is achieved by a variance stabilizing transform of the responses *r*, for which we use the Anscombe transformation, such that that transformed responses 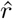 are obtained by

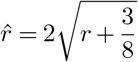

The SNR has been shown to serve well as a simple metric of data quality across diverse recording modalities and brain regions.

### Trajectories of spectral contrast analysis

We generated trajectories of spectral contrast tuning along the three behavioral dimensions (pupil size, instantaneous change of pupil size, and running speed). To that end, we generated 1000 MEIs per neuron, for all combinations of 10 values per behavioral parameter, for the 100 best predicted neurons of scan 1 in Fig. 2b. We sampled each behavioral parameters uniformly within the range of the 5th and 95th percentile. We then used the ARBTools toolbox (Walker et al., 2019b) to perform tricubic interpolation (Lekien & Marsden, 2005) of the 4D gridded data, given the 3 behavioral parameters and the spectral contrast value associated with each parameter combination. This allowed us to compute the gradients of spectral contrast at arbitrary locations within the range of behavioral parameters. To compute the trajectories, we started out from the behavioral state with the largest spectral contrast (i.e. highest green sensitivity). At that point, we computed the gradients with respect to the spectral contrast, and follow the gradient in the direction of minimal spectral contrast, with a step size of 0.05. We repeated this procedure until the magnitude of the gradient into the desired direction reached a cutoff value of 0.001. As this resulted in trajectories of different sample lenghts, we downsampled the trajectories by dividing them into *n* = 50 bins and computing the average in each bin. We used these downsampled trajectories to compute the average trajectories across the 100 neurons in 2h.

### Statistical analysis

We used Generalized Additive Models (GAMs) to analyze the relationship of MEI spectral contrast, cortical position and behavioral state (see Suppl. Statistical Analysis for details). GAMs extend the generalized linear model by allowing the linear predictors to depend on arbitrary smooth functions of the underlying variables (Wood, 2006). In practice, we used the *mgcv*-package for R to implement GAMs and perform statistical testing. For all other statistical tests, we used the Wilcoxon signed rank test for non-parametric data.

### Code and data availability

Our coding framework uses general tools like PyTorch, Numpy, scikit-image, matplotlib, seaborn, DataJoint, Jupyter, and Docker. We also used the following custom libraries and code: neuralpredictors (https://github.com/sinzlab/neuralpredictors) for torch-based custom functions for model implementation, nnfabrik (https://github.com/sinzlab/nnfabrik) for automatic model training pipelines using Data-Joint, nndichromacy for utilities, (https://github.com/sinzlab/nndichromacy), and mei (https://github.com/sinzlab/mei) for stimulus optimization. All data will be publicly available latest upon journal publication.

## Supplementary Information

**Supplemental Fig. 1.**
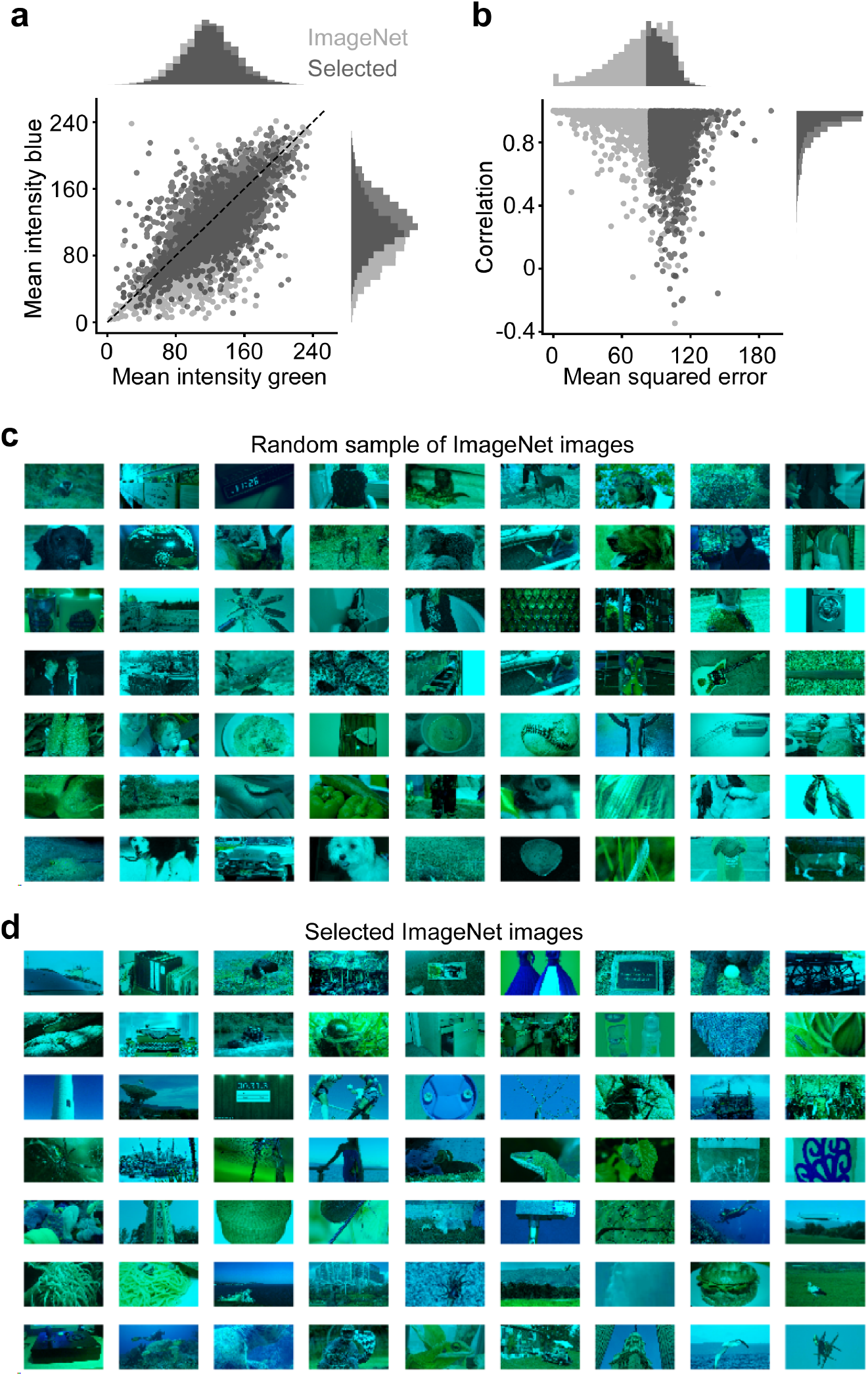
Selection of colored naturalistic scenes. **a**, Mean intensity in 8-bit pixel space of green and blue channel of randomly sampled ImageNet images (light gray; n=6.000) and selected images (dark gray; n=6.000; Methods). Images were selected such that the distribution of mean intensities of blue and green image channels were not significantly different. **b**, Distribution of correlation and mean squared error (MSE) across green and blue image channels (Methods). To increase chromatic content, only images with MSE > 85 were selected for visual stimulation. **c**, Random sample of ImageNet images. **d**, Sample of selected ImageNet images with high MSE.

**Supplemental Fig. 2.**
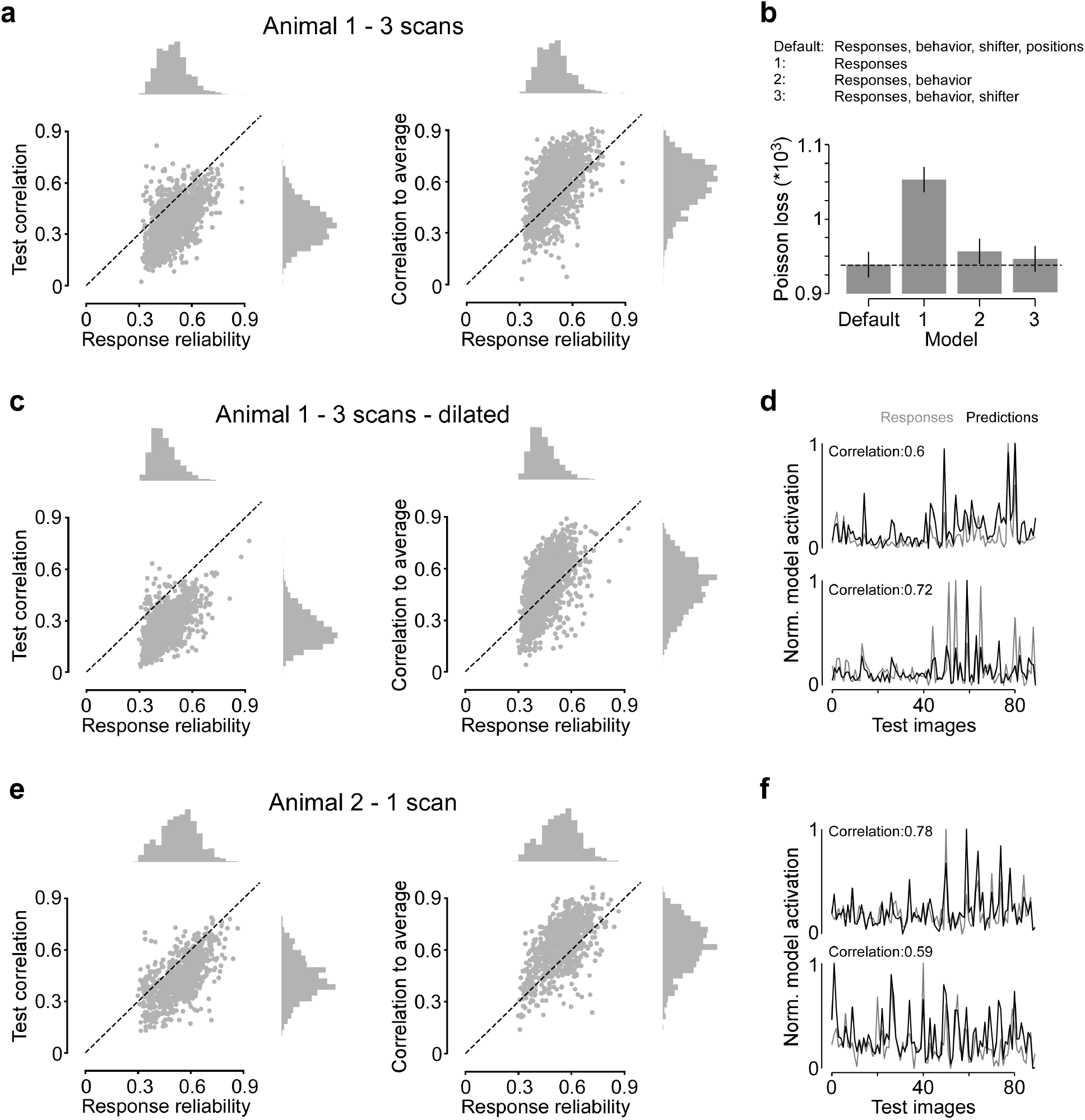
Model performance. **a**, Response reliability (Methods) plotted versus test correlation (left) and correlation to average (right) for data shown in Fig. 2 (n=1.759 cells, n=3 scans, n=1 mouse). **b**, Mean Poisson loss (lower is better) for different models trained on the dataset from (a). The default model is used for all analysis, while models 1-3 are shown for comparison. Dotted line marks mean Poisson loss of default model. The default model had significantly lower Poisson loss values compared to all three alternative models (Wilcoxon signed rank test: p<0.001, n=1,759). Error bars show 95% confidence interval. **c**, Like (a), but for scans with eye dilation shown in Fig. 4 (n=1,859 cells, n=3 scans, n=1 mouse). **d**, Mean responses to test images (gray) and model predictions (black) for two exemplary neurons from (c), with correlation coefficient of responses and predictions. **e, f**, Like (c, d), but for animal 2 shown in Fig. 6 (n=843 cells, n=1 scan, n=1 mouse).

**Supplemental Fig. 3.**
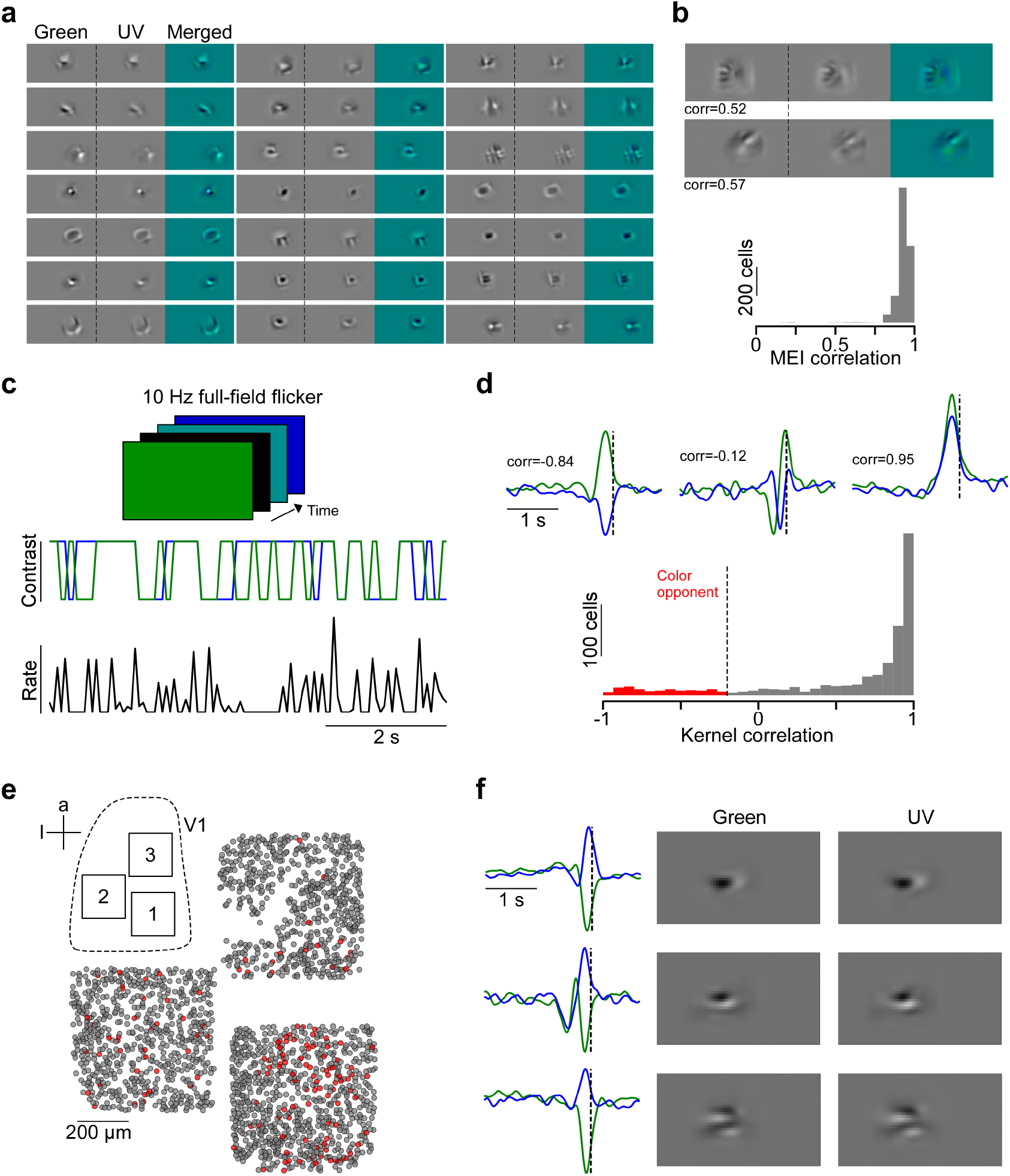
Spatial and temporal color opponency of mouse V1 neurons. **a**, MEIs of 21 exemplary neurons illustrate structural similarity across color channels. **b**, Distribution of correlation across color channels for dataset shown in Fig. 2. MEIs on top show example cells with relatively low correlation across color channels. **c**, Schematic illustrating paradigm of 10 Hz full-field binary white noise stimulus and corresponding response of exemplary neuron. **d**, Temporal kernels estimated from responses to full-field noise stimulus from (c) of three exemplary neurons (Methods) and distribution of kernel correlations (n=924 neurons, n=1 scan, n=1 mouse; scan 1 from (e)). Dotted line indicates correlation threshold of −0.25 – cells with a kernel correlation lower than this threshold were considered color-opponent (red). **e**, Neurons recorded in 3 consecutive scans at different positions within V1, color-coded based on color-opponency. **f**, Temporal kernels in response to full-field colored noise stimulus of three exemplary neurons (left) and MEIs of the same neurons. Neurons were anatomically matched across recordings by alignment to the same 3D stack (Methods).

**Supplemental Fig. 4.**
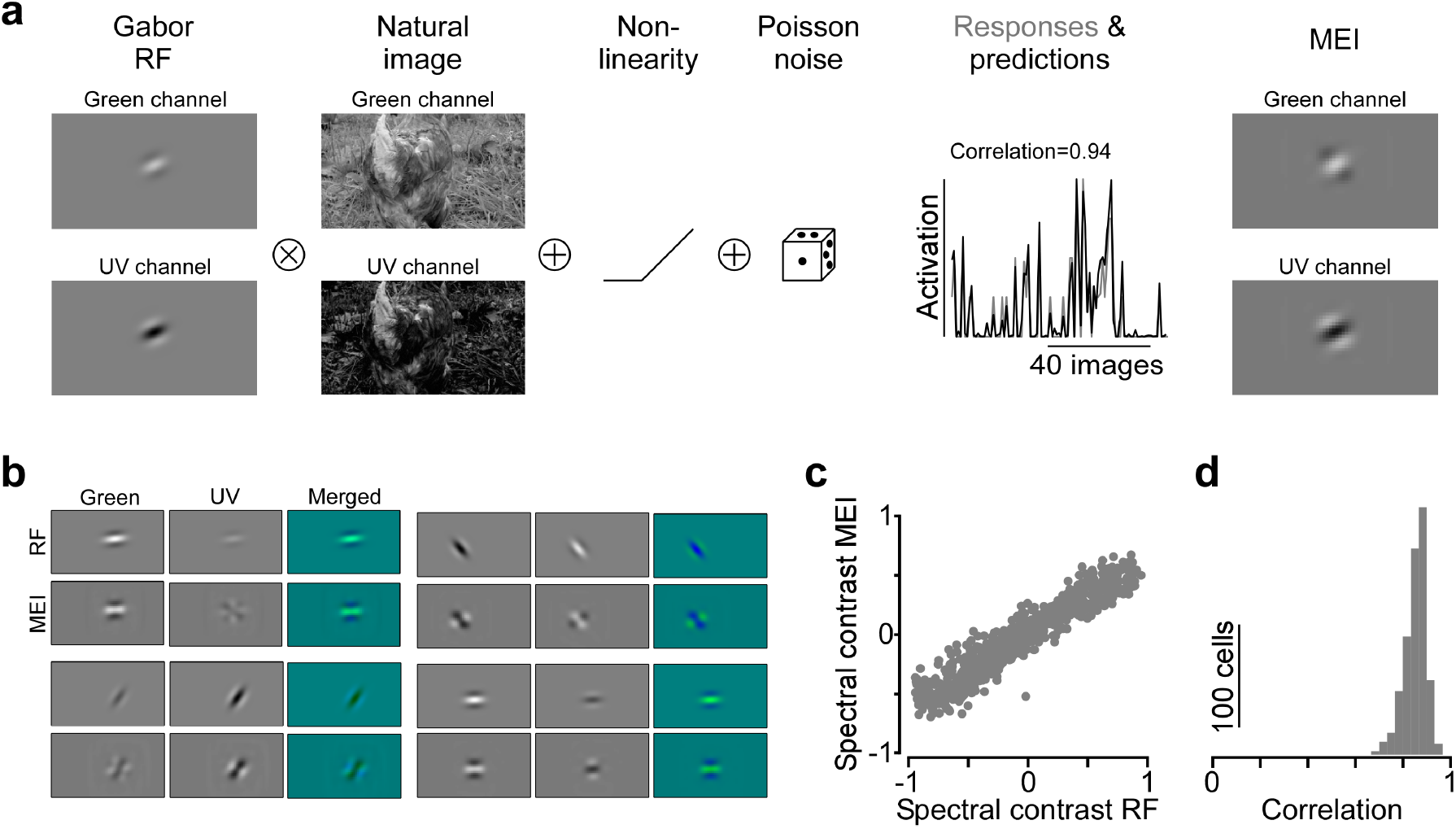
Model recovers color opponency and color preference of simulated neurons. **a**, We simulated neurons with Gabor receptive fields (RFs) of varying size, orientation, spectral contrast and color-opponency (correlation across color channels; Methods). Then, responses of simulated neurons with Gabor RFs were generated by multiplication of the RFs with the natural images also used during experiments. Corresponding responses were passed through a non-linearity and a poisson process before model training. Model predictions and optimized MEIs closely matched the simulated responses and Gabor RFs, respectively. **b**, Gabor RFs and corresponding MEIs of four example neurons, some of them with color-opponent RFs and MEIs. **c**, Spectral contrast of Gabor RF plotted versus spectral contrast of computed MEIs. The model faithfully recovered the simulated neurons’ color preference. Only extreme color preferences were slightly underestimated by our model, which is likely due to correlations across color channels of natural scenes. **d**, Correlation of the MEI with the ground truth gabor RF.

**Supplemental Fig. 5.**
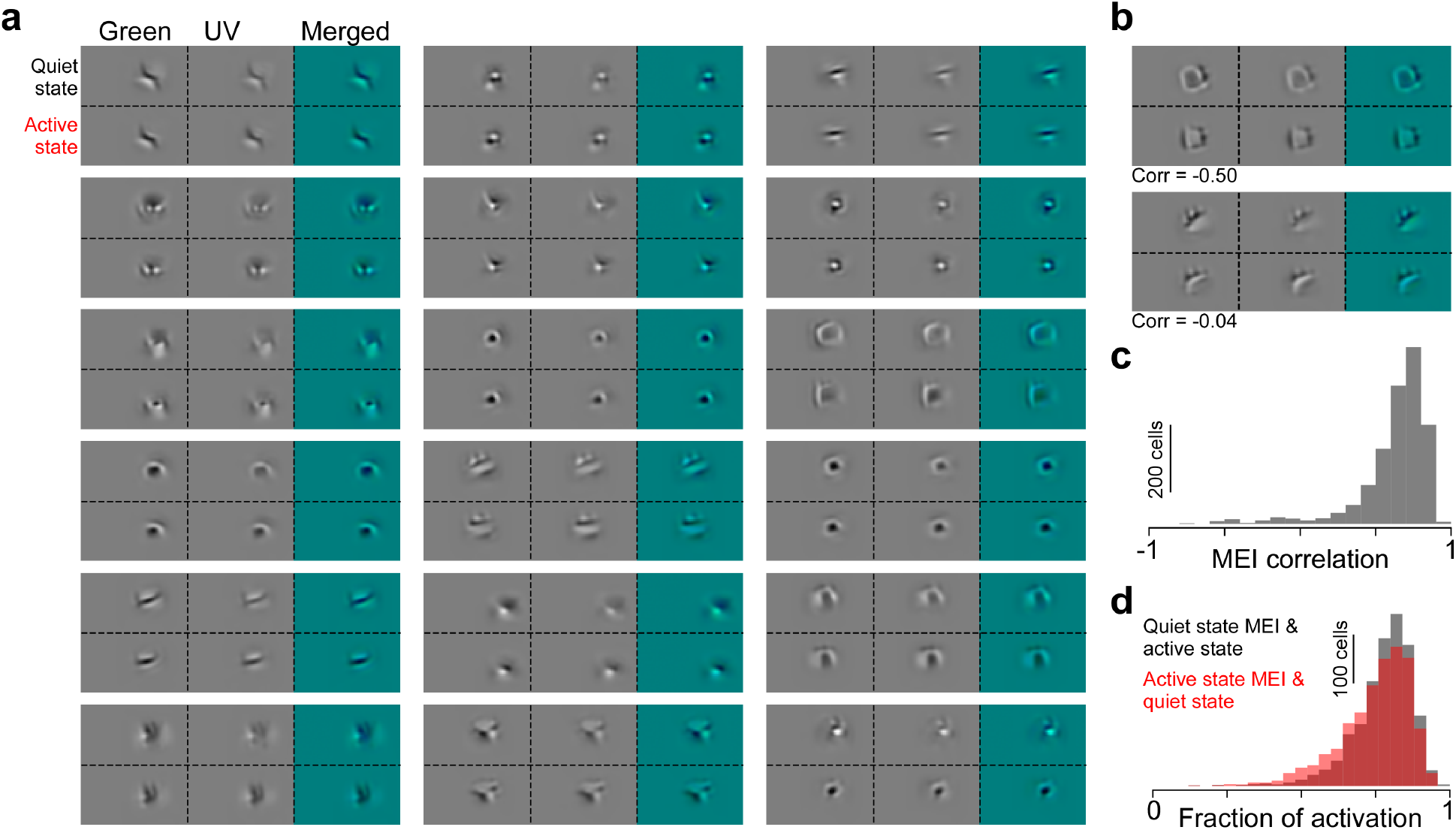
MEIs for quiet and active states. **a**, MEIs optimized for a quiet (top row of each sub-panel) and active (bottom row) behavioral state of 18 exemplary neurons illustrate structural similarity of MEIs across states. **b**, MEIs of two exemplary neurons with low correlation across behavioral states. **c**, Distribution of MEI correlation across states (n=1,759 neurons, n=3 scans, n=1 mouse). **d**, MEI activation in congruent behavioral state (n=1,759 neurons, n=3 scans, n=1 mouse). Gray: Model activation of MEI optimized for a quiet state presented to the model for active state relative to model activation of MEI optimized and presented for active state (activation=1). Red: Model activation of MEI optimized for active state presented to the model for quiet state relative to model activation of MEI optimized and presented for quiet state (activation=1).

**Supplemental Fig. 6.**
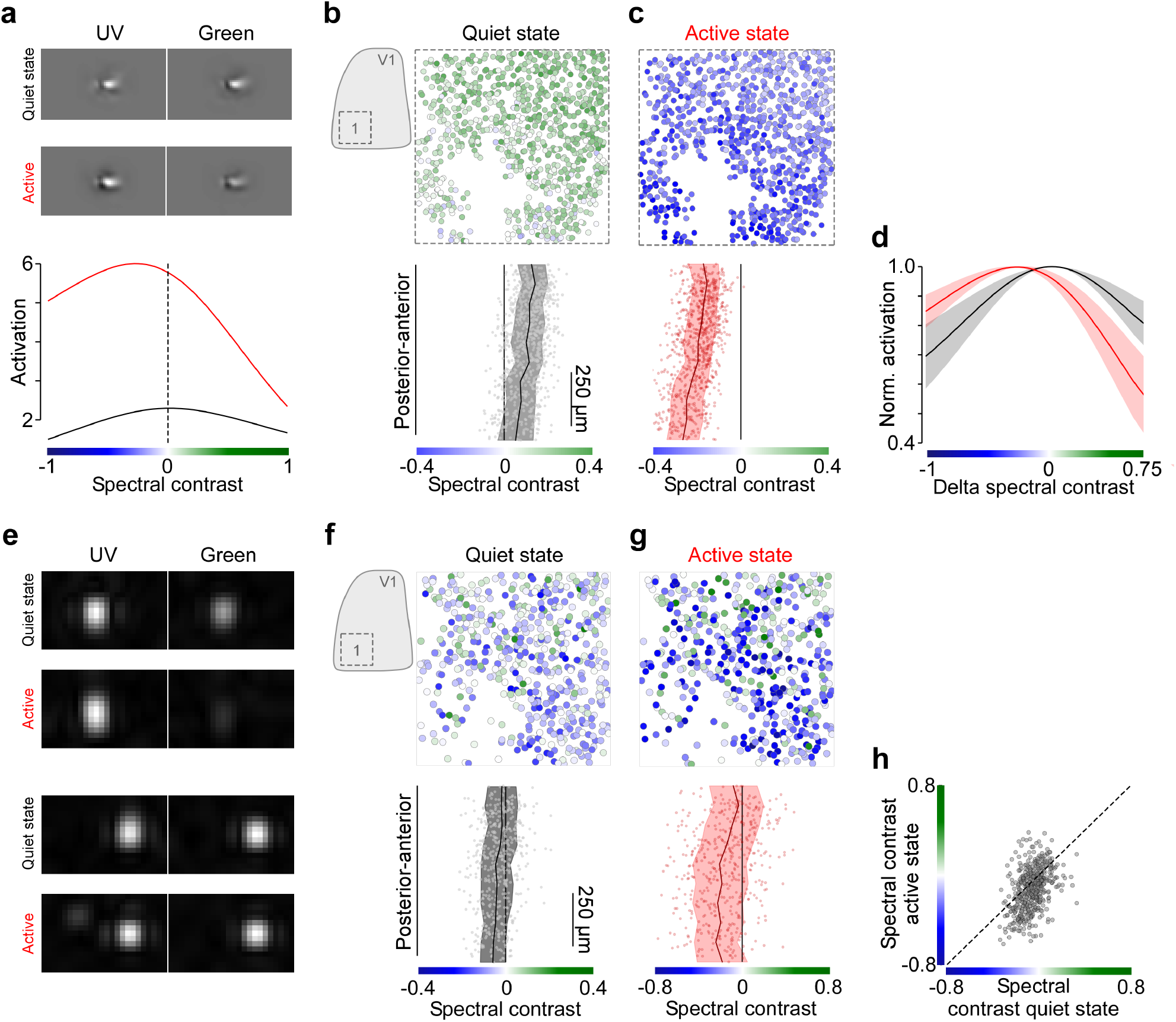
Behavioral modulation of color tuning of mouse V1 neurons - additional data. **a**, MEIs optimized for quiet and active state of exemplary neuron and corresponding color tuning curves. **b**, Neurons recorded in posterior V1 color coded based on spectral contrast of their quiet state MEI (top) and distribution of spectral contrast along posterior-anterior axis of V1. Black line corresponds to binned average (n=10 bins), with s.d. shading in gray. **c**, Like (b), but for active state. **d**, Mean of peak-normalized color tuning curves of neurons from (b, c), aligned with respect to peak position of quiet state tuning curves. **e**, Spike triggered average (STA) of two exemplary neurons estimated for small and large pupil trials. **f**, Neurons recorded in posterior V1 color coded based on spectral contrast of their small pupil STA (top) and distribution of spectral contrast along posterior-anterior axis of V1. Black line corresponds to binned average (n=10 bins), with s.d. shading in gray. **g**, Like (f), but for large pupil STA. **h**, Spectral contrast of small pupil STA plotted versus spectral contrast of large pupil STA.

**Supplemental Fig. 7.**
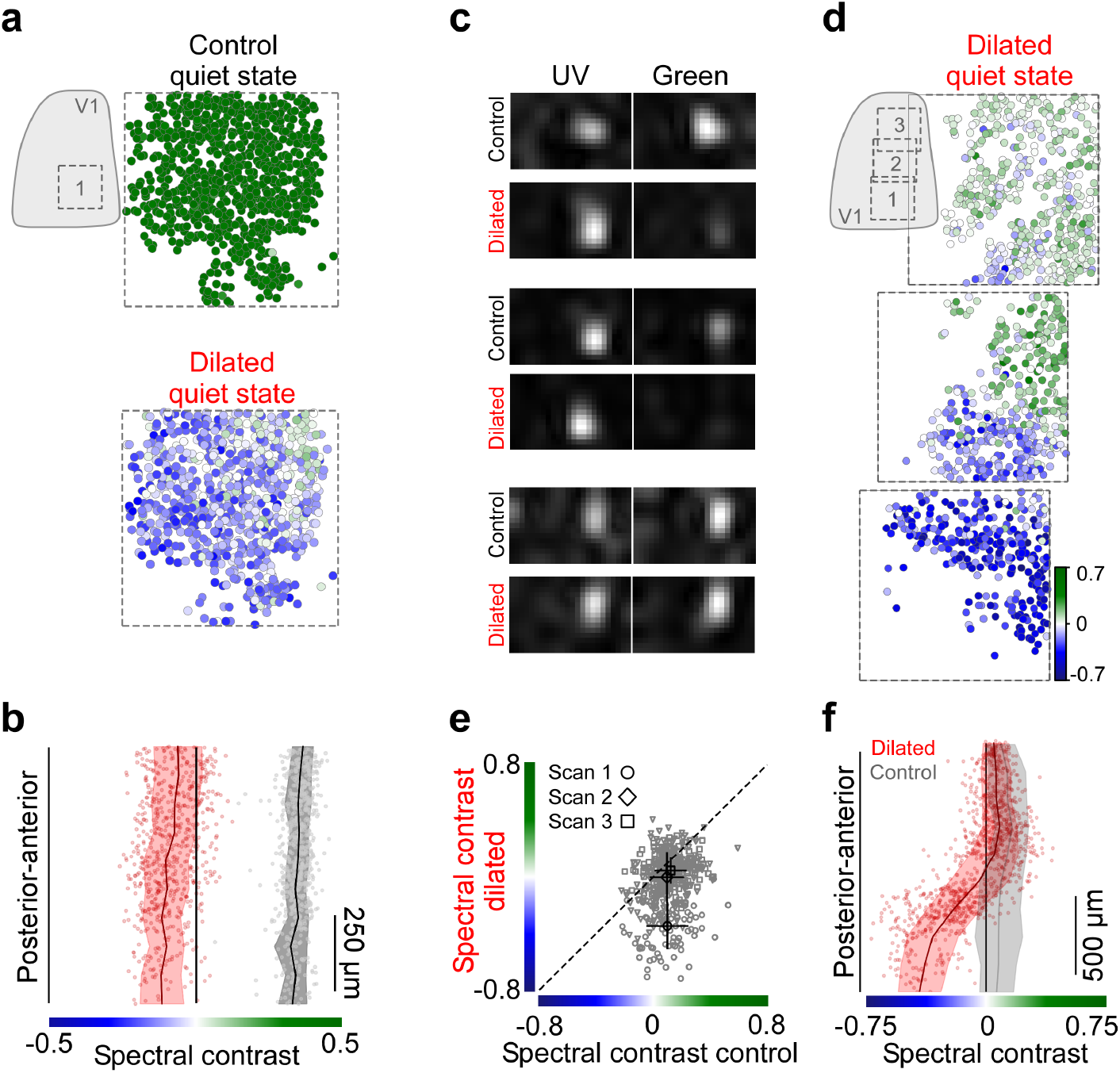
Pharmacological manipulations of pupil size. **a**, Neurons recorded in posterior-medial V1 color coded based on spectral contrast of their quiet state MEI under control condition (left) and for a scan with pupil dilation using atropine eye drops (right). **b**, Distribution of spectral contrast along posterior-anterior axis of V1 for control (gray) and dilated (red) conditions from (i). Solid lines correspond to binned average (n=10 bins), with s.d. shading. **c**, STAs of three exemplary neurons, estimated for quiet trials in control condition (black) and dilated condition (red). **d**, Neurons recorded in three consecutive experiments across the posterior-anterior axis of V1 (n=1,079 neurons, n=3 scans, n=1 mouse) color coded based on STA estimated for quiet trials in the dilated condition. **e**, Spectral contrast of quiet state STAs in control condition versus spectral contrast of quiet state STAs of anatomically matched neurons in dilated condition (n=444 neurons, n=3 scans, n=1 mouse). Mean and s.d. of each scan indicated in black. Wilcoxon signed rank test: p<0.001 (scan 1), p<0.01 (scan 2) and p<0.001 (scan 3). **f**, Spectral contrast of STAs of neurons from (f) along the posterior-anterior axis of V1 (red dots), with binned average (n=10 bins; red line) and s.d. shading. Black line and gray shading corresponds to binned average and s.d. of neurons recorded at the same cortical positions in control condition (cf. Fig. 3). Spectral contrast significantly varied across anterior-posterior axis of V1 for the dilated condition (n=1,079, p<0.001 for smooth term on cortical position of GAM; see Methods and Suppl. Statistical Analysis). Optimal spectral contrast changed with pupil dilation (n=1,079 (dilated) and n=943 (control), p<0.001 for condition coefficient of GAM; see Methods and Suppl. Statistical Analysis), with a significant interaction between cortical position and behavioral state modulation (see Suppl. Statistical Analysis).

**Supplemental Fig. 8.**
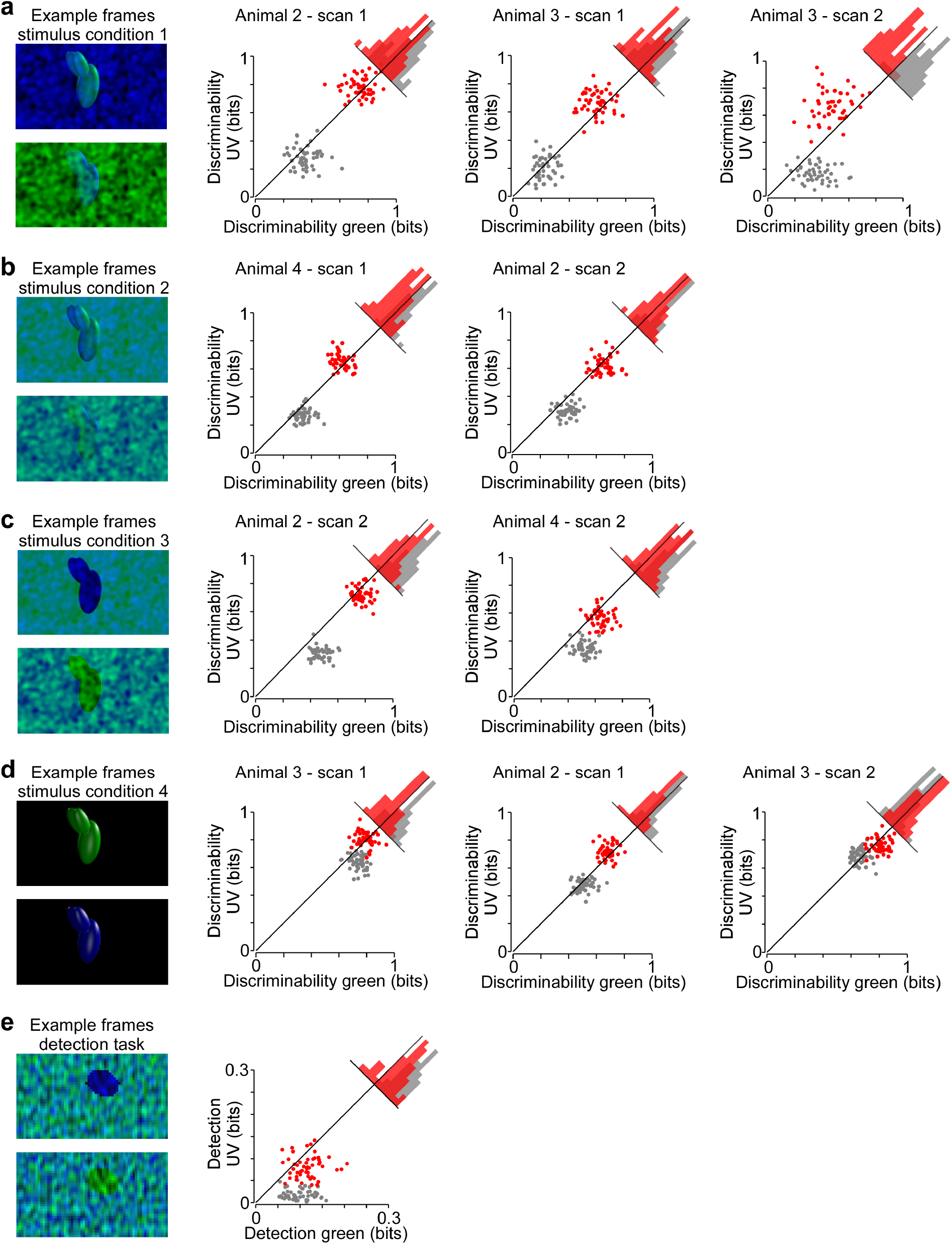
Additional data and stimulus conditions for decoding paradigm. **a**, Exemplary stimulus frame of condition 1 (shown in 7; green object with UV noise or UV object with green noise) and decoding results of three individual scans. Scatter plot shows discriminability of green versus UV objects for quiet (gray) and active (red) trials (Methods). Each dot represents a random sample of n=200 neurons of a neuron population (n=824) recorded in a single scan. Wilcoxon signed rank test (n=50): p=7.002e-06, p=8.707e-05, p=1.383e-09 **b**, Like (a), but for stimulus condition 2 with lower contrast of the object due to gray background in the object color channel. Wilcoxon signed rank test (n=50): p=1.725e-06, p=0.004 **c**, Like (a), but for stimulus condition 3 with objects as dark silhouettes and noise in the other color channel. Wilcoxon signed rank test: p=2.407e-05, p=0.0001 **d**, Like (a), but for stimulus condition 4 with high contrast objects and no noise in the other color channel. Wilcoxon signed rank test (n=50): p=3.108e-05, p=0.0004, p=0.0005 **e**, Exemplary stimulus frames of detection task from Fig. 7 and decoding result of an additional animal. Wilcoxon signed rank test (n=50): p=0.0005

## Supplemental Statistical Analysis – Generalized Additive Models

We used Generalized Additive Models (GAMs) to analyze the relationship of color preference, cortical position and behavioral state. GAMs extend the generalized linear model by allowing the linear predictors to depend on arbitrary smooth functions of the underlying variables (Wood, 2006):

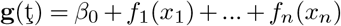

Here, *x_i_* are the predictor variables, *g* is a link function and the *f_i_* are smooth functions of the predictor variables. We used the *mgcv* package for R (version 1.8-24) to implement GAMs and perform statistical testing.

### MEI spectral contrast vs. cortical position across behavioral states

To model the dependence of MEI spectral contrast (`SC_MEI`) on cortical position (`Position`) and behavioral state (`State`), we used a Gaussian GAM with the factor behavioral state and separate smooth terms on cortical position for quiet and active states.

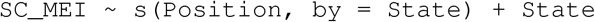

The basis dimension *k* and the smoothness was determined automatically by the `mgcv` package using generalized cross validation. The resulting model was fit using n=3,518 data points and yielded the following results:

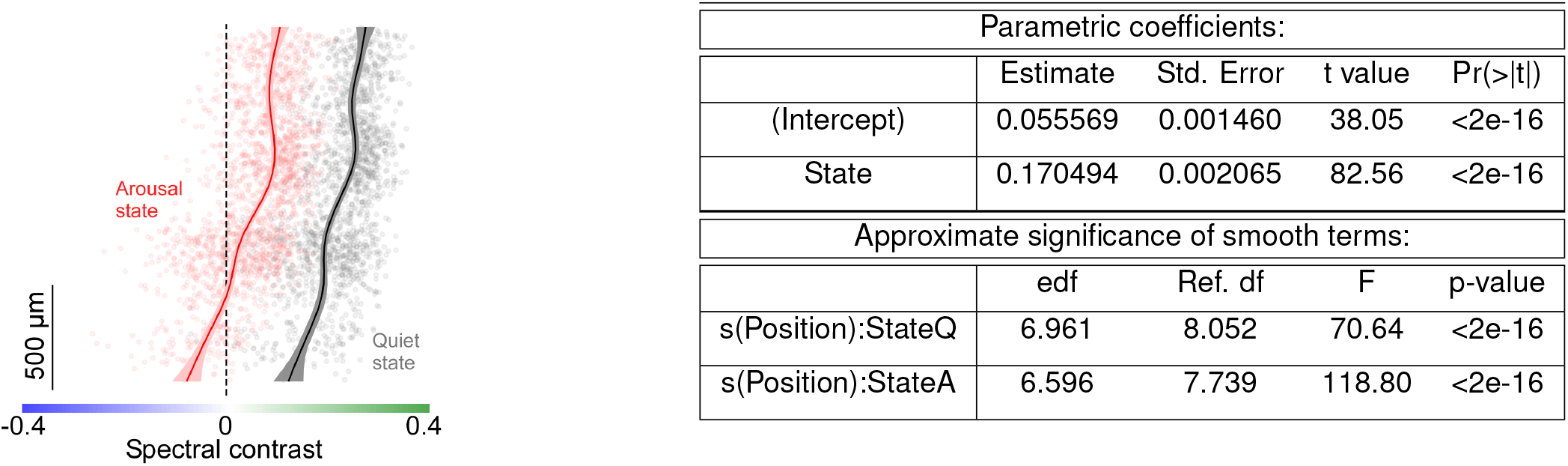

The (`State`) coefficient was highly significant, indicating that MEI color preference changes with behavioral state of the animal. In addition, the smooth terms for both quiet and active state were also highly significant, suggesting non-random variations of (`SC_MEI`) with cortical position.

Overall, the model explained 70.4% deviance. Using a GAM with only one smooth term for both behavioral states

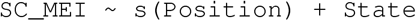

only slightly reduced the explained deviance (70.1%). This indicates that the dependence of (`SC_MEI`) on cortical position is similar for the quiet and active state tested here.

### STA spectral contrast vs. cortical position across behavioral states

To model the dependence of STA spectral contrast (`SC_STA`) on cortical position (`Position`) and behavioral state (`State`), we used a Gaussian GAM with the factor behavioral state and a smooth term for quiet and active states as a function of cortical position, as before.

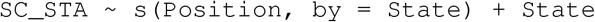

The basis dimension *k* and the smoothness was determined automatically by the `mgcv` package using generalized cross validation. The resulting model was fit using n=1,822 data points and yielded the following results:

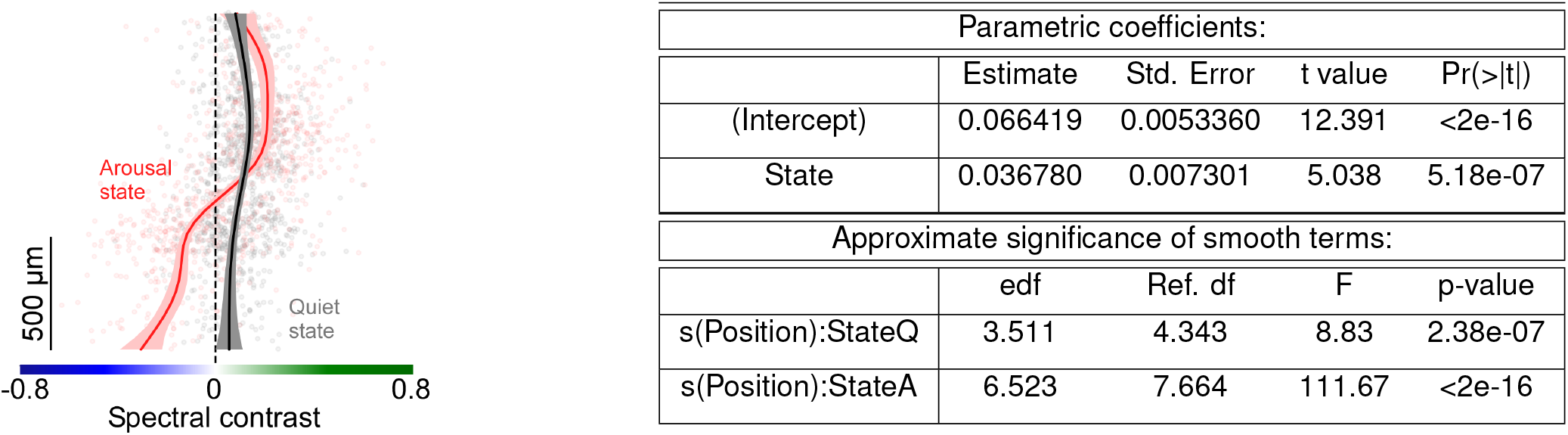

The (`State`) coefficient was highly significant, indicating that STA color preference changes with behavioral state of the animal. In addition, the smooth terms for both quiet and active state were highly significant, suggesting non-random variations of (`SC_STA`) with cortical position. (`SC_STA`) varied more strongly with cortical position for an active state, consistent with the lower p-value.

Overall, the model explained 34% deviance. Using a GAM with only one smooth term for both behavioral states

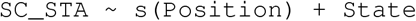

significantly reduced the explained deviance to 23%. This indicates that the dependence of (`SC_STA`) on cortical position differs across the two behavioral states.

### MEI spectral contrast vs. cortical position across control and dilated condition

To model the dependence of MEI spectral contrast (`SC_MEI`) on cortical position (`Position`) and control versus dilated condition (`Condition`), we used a Gaussian GAM with the factor behavioral state and a smooth term for control and dilated condition as a function of cortical position.

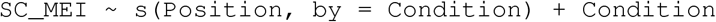

The basis dimension *k* and the smoothness was determined automatically by the `mgcv` package using generalized cross validation. The resulting model was fit using n=3,424 data points and yielded the following results:

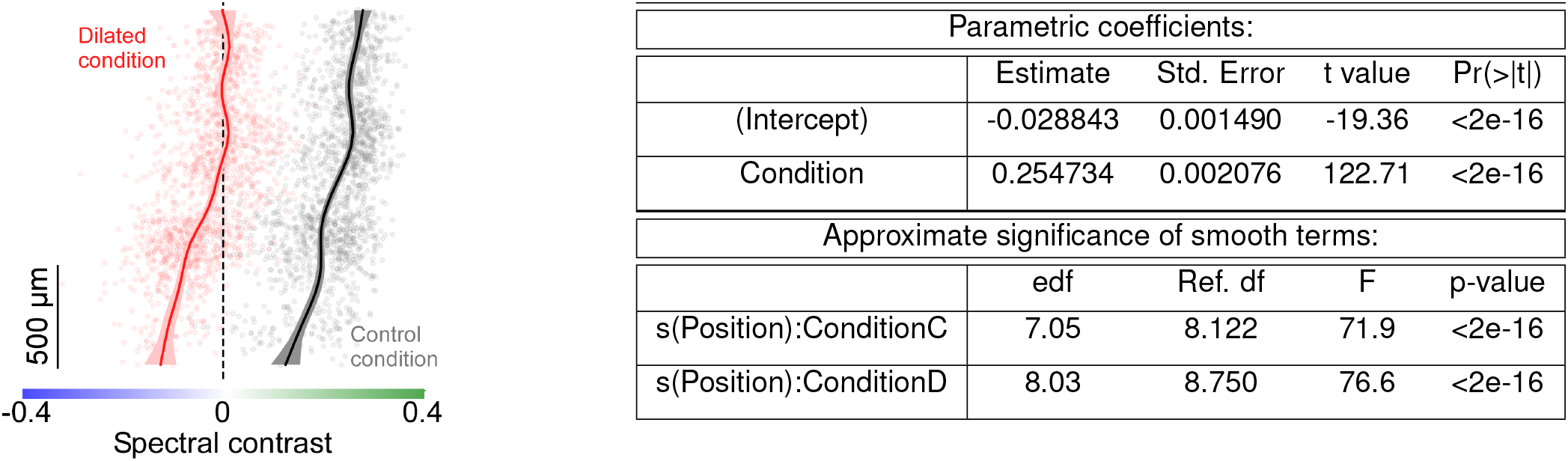

The (`Condition`) coefficient was highly significant, indicating that MEI color preference changes with pupil dilation. In addition, the smooth terms for both control and dilated condition were highly significant, suggesting non-random variations of (`SC_MEI`) with cortical position.

Overall, the model explained 82.9% deviance. Using a GAM with only one smooth term for both behavioral states

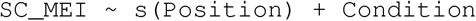

only slightly reduced the explained deviance (82.7%). This indicates that the dependence of (`SC_MEI`) on cortical position is similar for control and dilated condition.

### STA spectral contrast vs. cortical position across control and dilated condition

To model the dependence of STA spectral contrast (`SC_STA`) on cortical position (`Position`) and control versus dilated condition (`Condition`), we used a Gaussian GAM with the factor behavioral state and a smooth term for control and dilated condition as a function of cortical position.

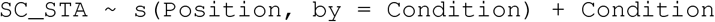

The basis dimension *k* and the smoothness was determined automatically by the `mgcv` package using generalized cross validation. The resulting model was fit using n=2,060 data points and yielded the following results:

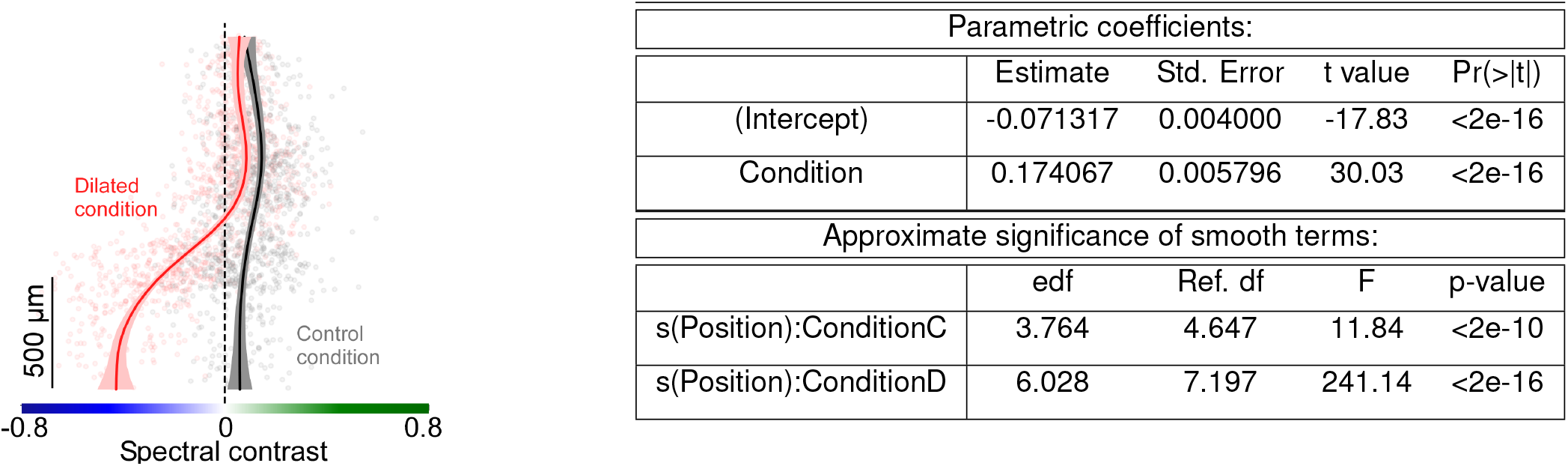

The (`Condition`) coefficient was highly significant, indicating that STA color preference changes with pupil dilation. In addition, the smooth terms for both control and dilated condition were highly significant, suggesting non-random variations of (`SC_STA`) with cortical position.

Overall, the model explained 57% deviance. Using a GAM with only one smooth term for both behavioral states

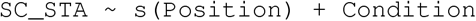

significantly reduced the explained deviance to 44.9%. This indicates that the dependence of (`SC_STA`) on cortical position differs across control and dilated condition.

